# DeepIFC: virtual fluorescent labeling of blood cells in imaging flow cytometry data with deep learning

**DOI:** 10.1101/2022.08.10.503433

**Authors:** Veera A. Timonen, Erja Kerkelä, Ulla Impola, Leena Penna, Jukka Partanen, Outi Kilpivaara, Mikko Arvas, Esa Pitkänen

## Abstract

Imaging flow cytometry (IFC) combines flow cytometry with microscopy, allowing rapid characterization of cellular and molecular properties via high-throughput single-cell fluorescent imaging. However, fluorescent labeling is costly and time-consuming. We present a computational method called DeepIFC based on the Inception U-Net neural network architecture, able to generate fluorescent marker images and learn morphological features from IFC brightfield and darkfield images. Furthermore, the DeepIFC workflow identifies cell types from the generated fluorescent images and visualizes the single-cell features generated in a 2D space. We demonstrate that rarer cell types are predicted well when a balanced data set is used to train the model, and the model is able to recognize red blood cells not seen during model training as a distinct entity. In summary, DeepIFC allows accurate cell reconstruction, typing and recognition of unseen cell types from brightfield and darkfield images via virtual fluorescent labeling.

## INTRODUCTION

Imaging flow cytometry (IFC) is a recent technique which combines fluorescent microscopy and flow cytometry into a high-throughput analysis platform (George *et al*. 2004). IFC allows for the study of cellular and molecular properties in fluidic samples at a single-cell level in high-throughput manner. It has been found useful in quantifying nucleic acids and protein expression (Doan *et al*. 2018), classifying rare cell types (Doan *et al*. 2018), identifying cells in their early apoptotic stage (George *et al*. 2004), examining host-intracellular parasites (Haridas *et al*. 2017) and other diagnostic purposes in hematology (Betters *et al*. 2015). In recent years, machine learning on IFC data (Luo *et al*. 2021) has been used to predict DNA content, quantify mitotic cell cycle phases (Blasi *et al*. 2016), reconstruct diabetic retinopathy disease progression and the cell cycle of Jurkat cells (Eulenberg *et al*. 2017) as well as to classify and identify white blood cells (Lippeveld *et al*. 2020, Nassar *et al*. 2019).

Fluorescent labeling requires a considerable amount of time, resources and effort, and can damage the cells (Icha *et al*. 2017). Consequently, so-called label-free or virtual staining approaches have been considered which may allow bypassing fluorescent labeling altogether. Recent studies in fluorescent imaging have used deep learning models to virtually stain brightfield images of adipose tissue (Wieslander *et al*. 2021), detect acute lymphoblastic leukemia cells (Doan, Case *et al*. 2020) and to discriminate between different cell lines (Matsuoka *et al*. 2021). Machine learning on IFC brightfield and darkfield images have been used to distinguish cell types (Lippeveld *et al*. 2020) and transitions between cell states (Eulenberg *et al*. 2017).

Label-free deep learning methods have been created to reconstruct fluorescent images from brightfield images (Christiansen et al. 2018, Ounkomol et al. 2018, Nguyen et al. 2021), but to our knowledge, the reconstruction of single-cell multichannel fluorescent images in IFC data has not been proposed. Label-free cytometry methods based on segmentation and unsupervised modeling (Nguyen et al. 2021), weak supervision (Otesteanu et al. 2021), cytometry by time of flight (CyTOF) (Hu et al. 2020) and time-stretch microscopy (Li et al. 2019) have been suggested. Previous methods have also utilized the Amnis IDEAS® software analyses, such as nuclear localization, or user generated cell masks (Lippeveld et al. 2020), which allow for the segmenting of the cell from its surroundings based on pixel intensity. Cell masks may negatively impact analysis, if their accuracy is not high enough (Dominical et al. 2017). They may also hinder recognizing the differences between cell states (Hennig et al. 2017). Spatial distribution of labels and dim-bright label continuum are also not explicitly modeled in these approaches.

In this study we present a novel method called DeepIFC (**Fig. 1)** to generate fluorescent images solely from morphological information in blood cell imaging flow cytometry data with minimal preprocessing. DeepIFC consists of a deep neural network model trained on brightfield and darkfield images of cells to generate corresponding fluorescent images. In contrast to many other approaches, DeepIFC reconstructs fluorescent images instead of predicting cell class labels such as the cell type, or fluorescent label intensity. The model also learns an intermediate representation of each input cell, which is useful in distinguishing cell types and features. We present tools to examine these representations (*i*.*e*., features) visually, and a method to identify cell types based on fluorescent images predicted by the DeepIFC model. Importantly, DeepIFC does not require manually annotated training data (*e*.*g*., cell type labels or cell image masks). We trained DeepIFC models on IFC data generated on peripheral blood mononuclear cells (PBMC), and evaluated the performance of these models also on data on red blood cells (RBC), a cell type not used in training. The DeepIFC workflow and models trained on PBMC data are available on GitHub (**https://github.com/timonenv/DeepIFC**).

**Figure 1.**
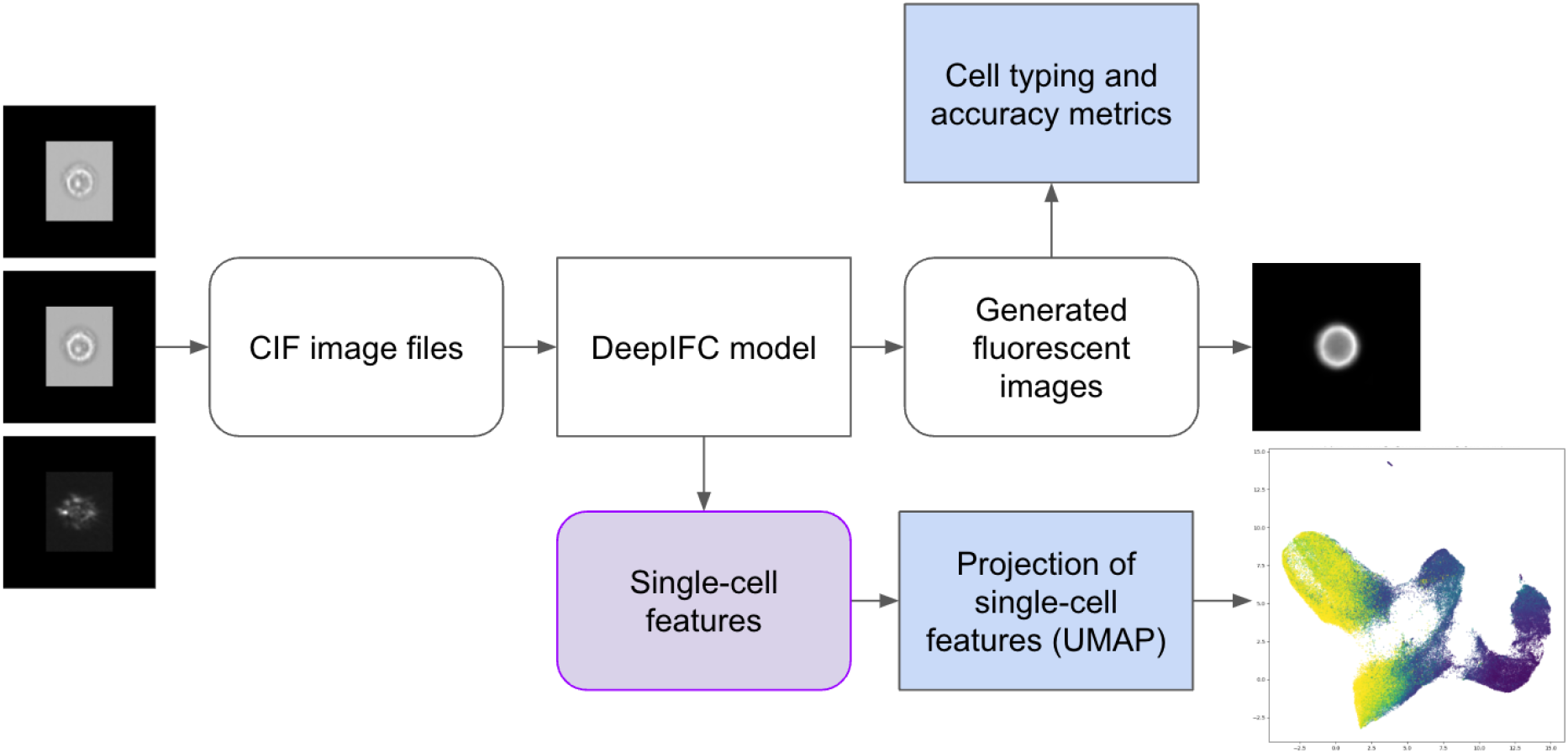
DeepIFC workflow. Imaging flow cytometry images are extracted from compensated image files (CIF) and image background intensities are normalized. Brightfield and darkfield images are processed by the DeepIFC model to generate corresponding fluorescence images. DeepIFC model also yields single-cell features, which the workflow visualizes by projecting the features onto a two-dimensional space with UMAP. The workflow also contains an interactive tool usable in a web browser for displaying the two-dimensional projection together with observed and generated images.

## RESULTS

### DeepIFC predictive performance and analysis of cellular features

DeepIFC models trained on PBMC IFC data (“complete dataset”, **Methods**) were found to accurately reconstruct fluorescent images of markers CD45 (*r*=0.90), CD14 (*r*=0.88) and CD3 (*r*=0.87), thresholding the average intensity values to determine marker positivity (CD45, AUROC=0.982; CD14, AUROC=0.979; CD3, AUROC=0.959) using only brightfield and darkfield images of the test dataset withheld from training (***Fig. 2a***). In addition to these surface markers, the reconstructions of images exhibiting positivity for 7-AAD were successful (*r*=0.79, AUROC=0.954). 7-AAD permeates the cell wall and binds to the DNA sequence of dead or damaged cells. On the other hand, fluorescent images for the surface markers CD56 (*r*=0.53, AUROC=0.805), CD19 (*r*=0.61, AUROC=0.800) and CD8 (*r*=0.41, AUROC=0.728) were less accurately reconstructed, likely due to the relatively small numbers of cells in the complete dataset exhibiting these markers (***Supplementary Table 1***) and morphological features unique to cells expressing surface markers such as CD8 and CD56 being difficult to distinguish in IFC images. Examples of measured images and images reconstructed by DeepIFC are shown in ***Supplementary Fig. 1* and *Supplementary Fig. 2***.

**Figure 2.**
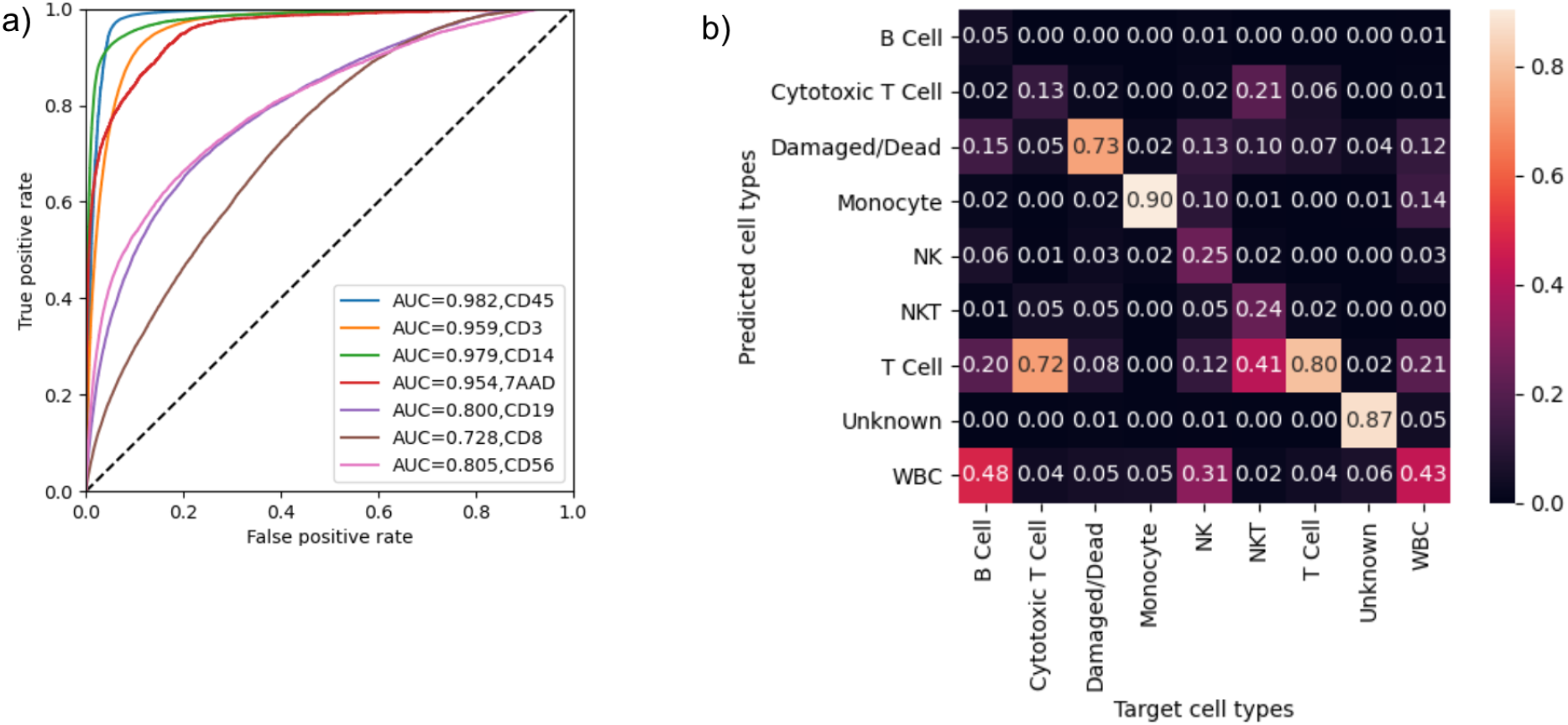
DeepIFC model predictive performance in PBMC data (“complete dataset”). **a**) Receiver operating characteristic (ROC) curve for each cell marker. **b**) Confusion matrix showing recall for the strict cell typing strategy where the ground truth (target cell types) is provided by observed fluorescent images (**Methods, Supplementary Fig. 4b**). Label “WBC” denotes a white blood cell only exhibiting the CD45 marker, and “Unknown” a cell exhibiting an unknown combination of markers not matching any cell type on the panel, or negativity for all markers.

To evaluate the ability of DeepIFC to identify cell types in the complete PBMC data, we assigned cell types based on thresholded fluorescent intensities in images generated by DeepIFC (strict cell typing strategy; **Methods**). We also performed the same cell typing procedure for observed images by manual gating in IDEAS*®* software (***Supplementary Fig. 4a***, gating strategy 1) as well as thresholding the mean intensity of each fluorescent image (***Supplementary Fig. 4b***) to establish two different ground truth settings. When compared to the image-based ground truth, DeepIFC was able to accurately classify monocytes (90% recall, 78% precision) likely due to their distinct morphology (***Fig. 2b, Supplementary Table 3***). Despite both CD3 and CD45 markers being well predicted individually, performance predicting CD8- CD56- T cells was found to be at the moderate level (80% recall, 73% precision). This was mostly due to falsely predicted CD8 and 7-AAD label fluorescence, since 6% and 7% of CD8- CD56- T cells were predicted to be cytotoxic T cells, or dead or damaged cells, respectively. Dead or damaged cells were predicted at 73% recall and 54% precision. DeepIFC showed high performance of 87% recall and 92% precision when predicting cells of unknown type, that is, cells where the predicted marker fluorescences did not correspond to any known combination or are all negative (**Fig. 2b, Supplementary Fig. 4b, c**). In contrast to these well-predicted types, the cell types appearing in smaller amounts in the data or exhibiting T cell subtype markers were predicted at much lower recall levels (NK, 25%; NKT, 24%; cytotoxic T, 13%; B, 5%). Unsurprisingly, the most common incorrect prediction for NKT cells was a T cell, with 41% of true NKT cells classified as T cells. Likewise, 72% of true cytotoxic T cells were classified as T cells. Of true NK cells, 31% were identified as WBCs due to failure to predict CD56 fluorescence from morphology. The predictive performance on multiple cell types improved when dead or damaged, unknown cells and cells positive for only the CD45 marker were removed from analysis. Most notably, recall and precision for monocytes improved from 90% and 78% to 98% and 92%, respectively (***Supplementary Table 3***). Cell type fractions predicted by DeepIFC were found to correspond well to fractions obtained from observed images and analysis in IDEAS*®* (**Figure 3, Supplementary Table 1**).

**Figure 3.**
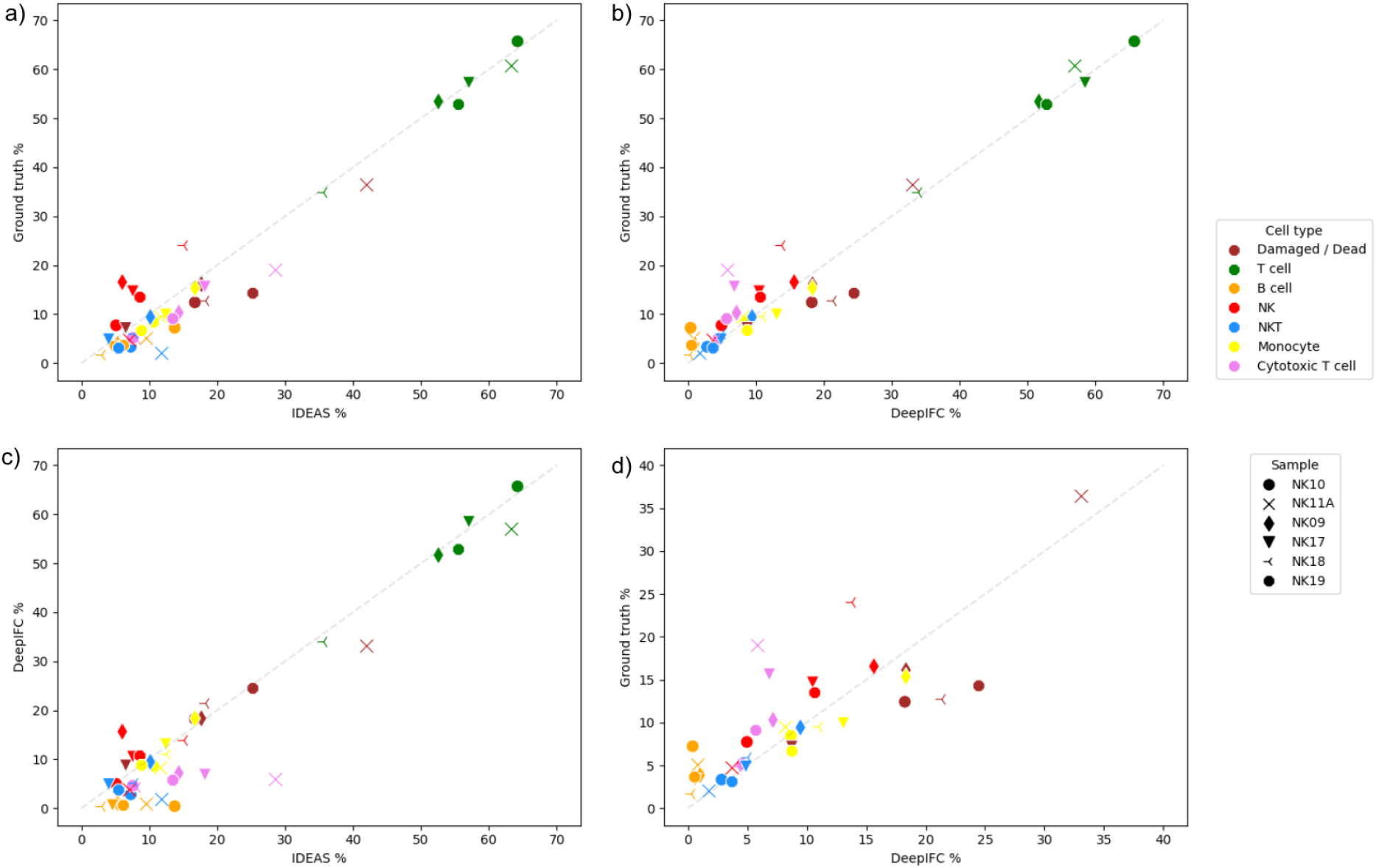
Cell type fractions obtained from gating strategy 1 performed in IDEAS®, and thresholding of average fluorescence intensities performed for the ground truth images and DeepIFC predicted images as per the permissible cell typing strategy, for each donor sample. Number of cells indicated for each sample. Comparison of **a**) image-based ground truth and IDEAS® cell type fractions, **b**) image-based ground truth and DeepIFC generated cell type fractions, and **c**) DeepIFC generated cell type fractions and IDEAS® cell type fractions. **d**) An inset of the image-based ground truth and DeepIFC comparison restricted to range [0, 40%].

We then extracted features learnt by the DeepIFC model for each cell in the complete PBMC dataset, and visualized them by projecting the features to two dimensions with UMAP (McInnes *et al*. 2018). We observed four distinct large clusters (***Fig. 4***) corresponding to T cells (cluster 1), monocytes (2), dead or damaged cells (3), and objects which did not exhibit any fluorescent label or had unknown combinations of markers (4). In addition, two smaller clusters consisting of debris were visible (5a, 5b). A subset of cells (n=5778; 1%) expressing both the monocyte marker CD14 and dead/damaged marker 7-AAD were found to connect the monocyte and dead/damaged cell clusters. These dead/damaged monocytes are most likely morphologically distinct enough from other cells so that they form a bridge between the two clusters instead of mixing with other dead/damaged cell types (3). Although rarer cell types, that is, cell types exhibiting T cell subtype markers in the data did not constitute separate clusters, we found NKT cells to be concentrated to the bottom left of cluster 1, while NK and B cells were found predominantly in the bottom right of cluster 1. Similar clustering of cytotoxic T cells was not found.

**Figure 4.**
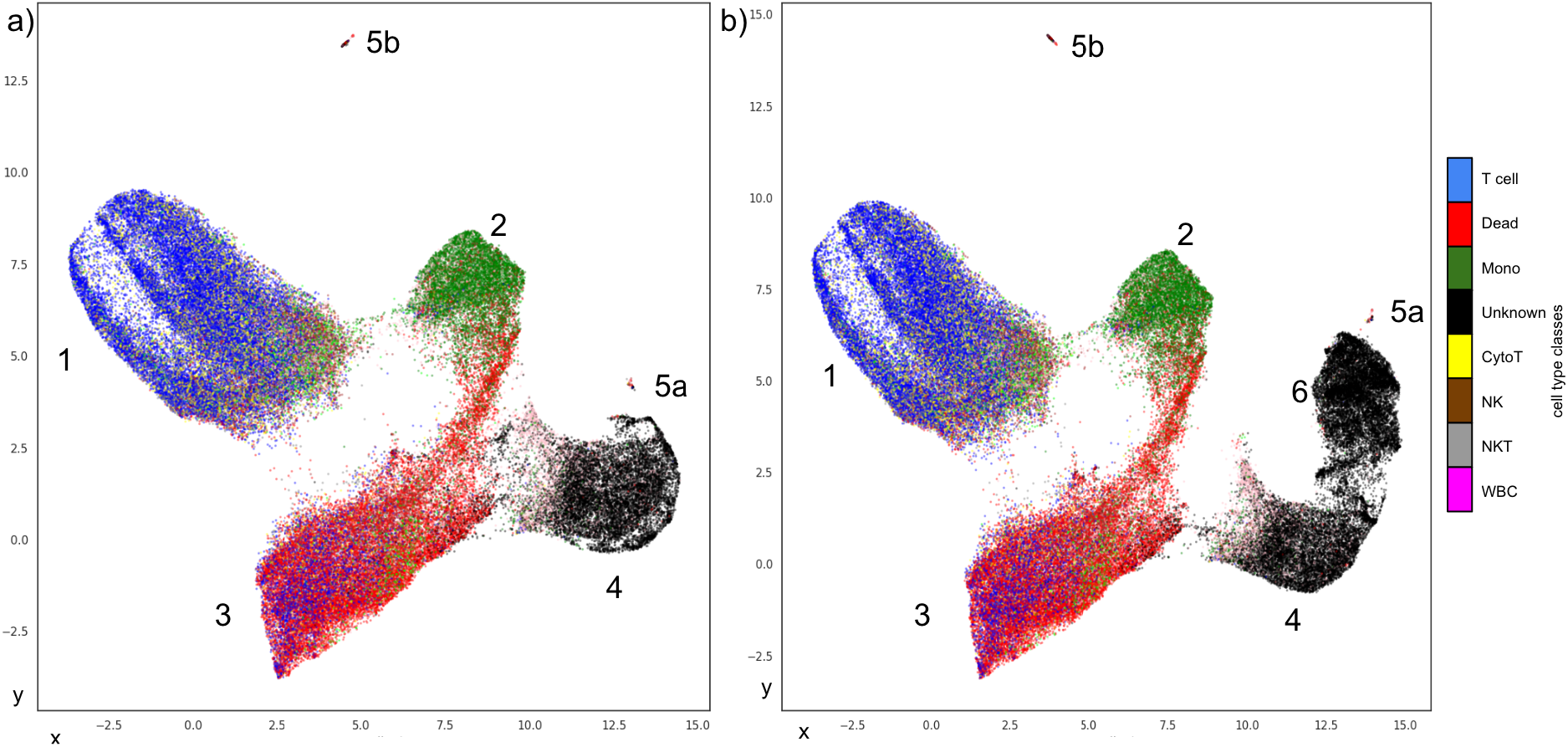
UMAP projections of single-cell features learnt by DeepIFC. Features were extracted and combined from the seven DeepIFC models learnt for each cell type in the data. **a**) Data from the DeepIFC models trained on complete PBMC data. Cell types identified with the strict typing strategy according to observed images are indicated with colors. Four main clusters are visible: T cells (1), monocytes (2), dead or damaged cells (3), and objects not expressing any fluorescent marker or otherwise unknown (4). Two smaller clusters containing debris are indicated with 5a and 5b. **b**) UMAP projection showing combined PBMC and RBC data (Doan et al. 2020). DeepIFC distinguishes RBCs as a distinct entity (cluster 6) based on their morphology despite not seeing large numbers of these types of cells during model training. As red blood cells do not express any marker, they are correctly identified as “unknown” class along with other cells negative for all markers, or with unknown combinations of positive markers. The RBC cluster also connects to the unknown/all negative cluster, but forms a distinct entity.

### Prediction of doublet events

The PBMC IFC data contained a number of acquisition events where two (doublets) or more cells were imaged at the same time. Scrutiny of fluorescent images generated by the DeepIFC complete data model revealed that the method was able to predict label fluorescence separately for multiple cells in the same event (***Supplementary Fig. 3***). We found the doublet events to contain proportionally fewer T and cytotoxic T cells compared to all events (T, 58.9% of cells in doublet events, 95% CI 54.9–62.7% vs 65% of cells in all events; cytotoxic T, 10.5%, 95% CI 8.2–13.1% vs 15%), while exhibiting a two-fold increase in monocytes (24.8%, 95% CI 21.5–28.3% vs 12%). T cell - T cell doublets were the most common at 23% of all doublet events. Other cell doublets were more rare, e.g. cytotoxic T cell - T cell pairing (13% of all doublet events), NKT - T cell (6%), monocyte - T cell (6%) and natural killer - T cell (6%). Monocytes were most often paired with other monocytes (18% out of all doublet events).

### DeepIFC recognizes a cell type not seen during training

To understand whether DeepIFC models would be useful in analyzing cell types not seen during training, we processed brightfield and darkfield images of red blood cells from a recent IFC study (Doan *et al*. 2020; dataset Mixed_CE47_D2) with the DeepIFC model trained on PBMC images (“complete dataset”) without retraining the model on RBC images, and computed the DeepIFC features for the RBC images. Surprisingly, a large fraction of these RBCs (97%) formed a new cluster (Cluster 6, **Fig. 4b**) separate from the mononuclear cell clusters, demonstrating the ability of DeepIFC to recognize cells with unseen morphology as a distinct entity.

### Balancing training data improves DeepIFC prediction performance

Finally, we investigated whether it would be possible to improve DeepIFC performance on cell types which were poorly predicted in the complete PBMC dataset. To do this, we created balanced datasets separately for each cell type such that the proportion of the target cell type was set to 50% (**Methods**). DeepIFC models were trained on multiple balanced datasets with different amounts of target cells to study the effect of number of cells on performance. Prediction performance of DeepIFC models showed substantial improvements on multiple cell types over the performance of models trained with the unbalanced, complete dataset (***Fig. 5, Supplementary Fig. 5, Supplementary Table 3***). For most cell types, the saturation point in performance increase was reached at 3200 cells, with the rise in accuracy slowing down with larger cell amounts. Notably, a large performance gain was observed with NKT cells, which were predicted at 83% recall and 63% precision in balanced data compared to 24% recall and 21% precision in unbalanced data. Similarly, recall of NK, cytotoxic T and B cells increased from 25% to 67%, 13% to 68% and 5% to 54%, respectively. Results for average accuracies over three training runs for each model are shown in **Figure 5**, and the results for best performing individual models are shown in **Supplementary Figure 5** and **Supplementary Table 3**.

**Figure 5.**
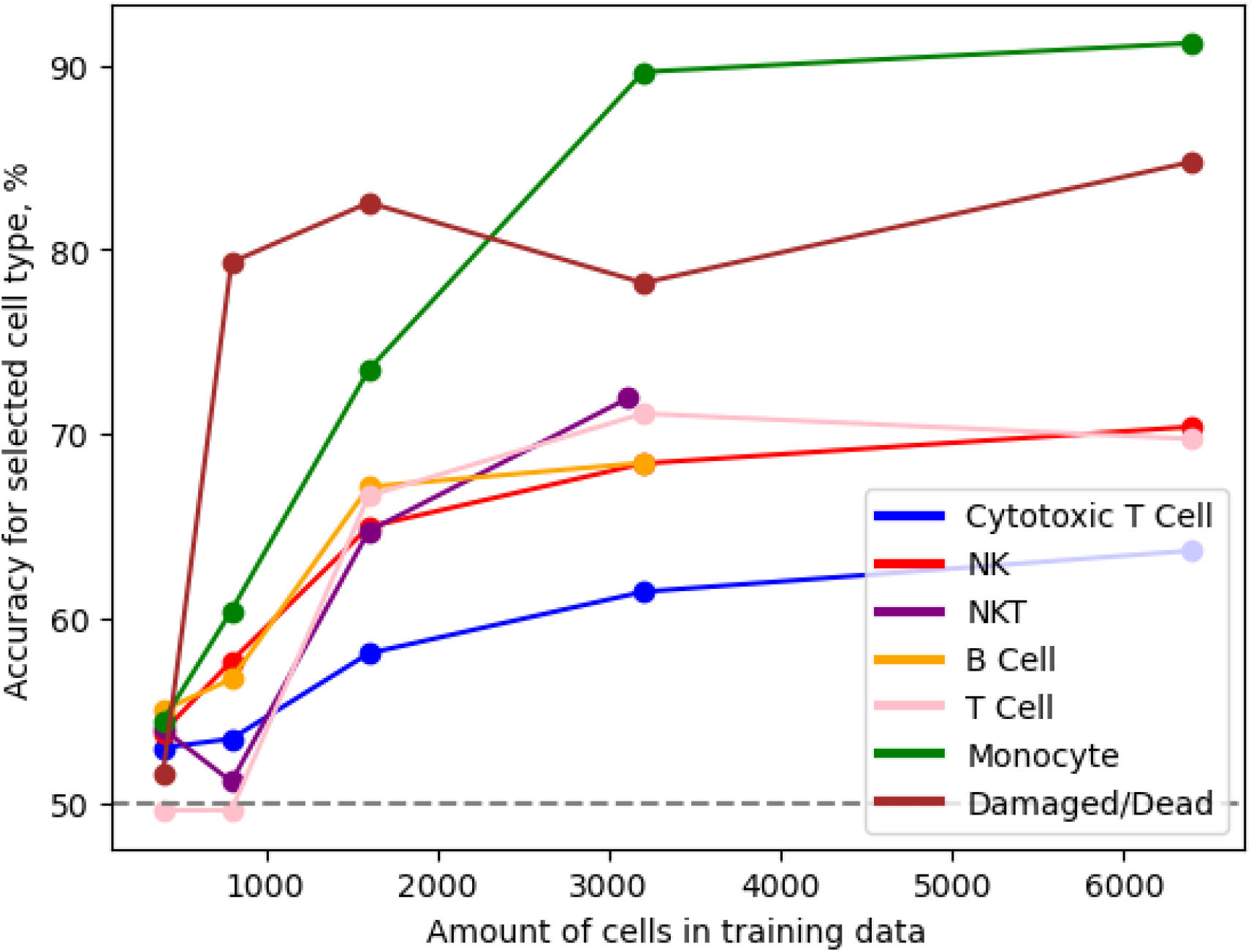
DeepIFC binary prediction accuracy (Y-axis) in the balanced PBMC datasets with respect to the number of cells of the target cell type (X-axis). Target cell type indicated by the color. Average accuracies over three replicates (i.e., training runs) are shown.

## DISCUSSION

In this study, we presented a novel computational method called DeepIFC for reconstructing fluorescent images from brightfield and darkfield images acquired with an imaging flow cytometer. Although generating images across multiple microscopy modalities has been demonstrated in fluorescence microscopy (Wang *et al*. 2019), to our knowledge DeepIFC is the first method employing such cross-modality learning in multichannel IFC. DeepIFC trained on IFC data from PBMCs showed high accuracy in reconstructing fluorescent images for the cell surface markers CD45, CD3 and CD14 solely from morphology. The method was additionally able to accurately recognize dead and damaged cells, which may enable sample quality control in IFC without explicitly staining for damaged cells thus allowing a fluorescent channel to be used for other purposes (George *et al*. 2004). Interestingly, no distinct difference in dead or damaged cell morphology from live ones was visible to the expert eye in brightfield and darkfield images, despite DeepIFC predicting the fluorescence of dead/damaged cell marker 7-AAD reasonably well. This may be due to the phase of apoptosis where no changes to cell morphology clearly visible to human experts have occurred yet, but the cells already bind 7-AAD. Accurately predicting quality of cells is of critical importance in operation of blood services and biobanks, and thus models such as DeepIFC hold promise to decrease costs and improve throughput in these facilities.

DeepIFC enables identification of cell types and characteristics via virtual gating of the generated fluorescent images. DeepIFC models achieved classification performances ranging from 54% (B cells) to 92% (monocytes) for recall, and from 61% (cytotoxic T cells) to 91% (monocytes) for precision. The reason for the accurate prediction of certain cell types (CD45+ leukocytes, CD3+ T cells, CD14+ monocytes, 7-AAD+ dead cells) may be that they were either present in high amounts in the data (leukocytes, T cells) or their morphology was distinct from the other cell types (monocytes, dead cells). Cell types appearing in smaller amounts were predicted at much lower recall levels in the complete, unbalanced dataset (NK, 25%; NKT, 24%; cytotoxic T, 13%; B, 5%), however training with balanced data resulted in substantially improved prediction performance compared to unbalanced original data in most cell types (NK, 67%; NKT, 83%; cytotoxic T, 68%; B, 54%) except for CD8- CD56- T cells. Similarly to previous efforts utilizing machine learning (Lippeveld *et al*. 2020), DeepIFC had difficulties distinguishing between T cell subtypes. Morphological differences between T cell subtypes (CD56, CD8) were not found to be visible to the human eye in brightfield images. Regardless, we found DeepIFC to classify the subtypes better than random guess (cytotoxic T, 64% binary classification accuracy; NKT, 70%).

A unique feature of our approach compared to previous IFC data analysis methods which attempt to predict marker positivity as labels (Eulenberg *et al*. 2017, Nassar *et al*. 2019, Lippeveld *et al*. 2020) is that DeepIFC reconstructs the entire fluorescent image instead of outputting a binary or class-based prediction. This allows the method to consider the spatial distribution of fluorescence as well as predict fluorescence in multiple cells in the same IFC acquisition event, and offer the user visual cues on the virtual fluorescent labeling process. These capabilities may enable analysis of complex cell-cell interactions without fluorescent labels (Burel *et al*. 2020). To this end, DeepIFC was able to identify different cell types in doublet events, highlighting a two-fold increase in the proportion of monocytes in doublet events (25% in doublets vs 12% in all events). Monocyte-monocyte doublets are found in e.g. psoriasis patients’ blood (Golden *et al*. 2015), while T cells are known to form doublets with monocytes in the event of infection (Burel *et al*. 2019).

We demonstrated how DeepIFC models can be applied to data not seen during training, for example from cell types not present in training data, to detect novel cell entities. Analyzed with a DeepIFC model trained on data from PBMC samples with RBCs removed, a set of RBCs from an independent study (Doan, Sebastian *et al*. 2020) formed a distinct cluster from mononuclear cell types. Machine learning methods able to distinguish labels not present in training datasets (*i*.*e*., zero-shot learning) have been studied extensively (Wang *et al*. 2019). In this study we showed for the first time these capabilities applied to IFC. We envision DeepIFC models trained on larger datasets to be able to distinguish a wide variety of cell types and cellular characteristics. As RBCs have morphology distinct from mononuclear cells, it would be interesting to explore DeepIFC’s performance also on other cell types not present in the datasets used in this study.

In the future the effect of larger, balanced and augmented datasets may be examined to improve the performance of DeepIFC, as suggested by Lippeveld *et al*. (2020). Cell masks such as those generated by IDEAS*®* may reduce the possible bleedthrough between channels. The quality of the samples may also play a role. The samples utilized here were frozen and thawed, and there was substantial variation in the amount of dead cells per sample (6.6%–42.0%). Suboptimal cryopreservation may cause alterations of the cellular phenotype and lead to non-specific binding of antibodies (Germann *et al*. 2013, Tomlinson *et al*. 2013) as well as changes in morphology. On the other hand, analysis of fresh samples, especially from patients, is not technically possible. The method should, thus, be able to handle suboptimal samples. Other improvement could be further optimization of the antibody panel by testing different fluorochromes for different markers.

Taken together, methods such as DeepIFC able to perform virtual labeling and cell type identification solely from morphology hold promise to transform diagnosis of hematological diseases and blood processing pipelines by not having to introduce fluorescent labels during workflow, reducing costs and processing time required. Possible avenues to develop the method further include utilizing larger training datasets covering more cell types, data augmentation to improve performance on rare cell types (Luo, Nguyen *et al*. 2021), and a user-friendly graphical tool to use the software.

## CODE AVAILABILITY

The DeepIFC workflow and models trained on PBMC data are available under a permissible license. An example of the interactive UMAP tool for a subset of the complete MNC data can be found here: **https://timonenv.github.io/DeepIFC/**. The code, trained models and requirements for the DeepIFC workflow are found here: **https://github.com/timonenv/DeepIFC**.

## ACKNOWLEDGEMENTS

This study was supported by grants from the Academy of Finland (322675, 328890). We would like to thank CSC – IT Center for Science, Finland, for generous computational resources. We thank Tomi Määttä for valuable comments on the manuscript, and Lotta Andersson, Prima Sanjaya, Loïc Masson, Olle Hansson, Jarno Laitinen and Timo Miettinen for technical assistance.

## METHODS

### DeepIFC model

We developed a computational workflow called DeepIFC to perform virtual fluorescent labeling on brightfield and darkfield IFC images (**Fig. 1**). For each input cell, DeepIFC workflow results in a generated image for each fluorescent channel, and a set of single-cell features. The generated images are then used to perform virtual gating to classify cell types. DeepIFC also visualizes the single-cell features by projecting the features onto a two-dimensional space with Uniform Manifold Approximation and Projection for Dimension Reduction (UMAP) (McInnes *et al*. 2018).

At the core of DeepIFC is a deep neural network model based on the Inception U-Net architecture (Cahall *et al*. 2019) (***Fig. 6***). Inception U-Net combines the U-Net architecture common in image segmentation tasks (Ronneberger *et al*. 2015) with Inception modules (Szegedy *et al*. 2015). The model processes input images through consecutive Inception modules, where the spatial dimensions are first contracted towards the model bottleneck (***Fig. 6***, contracting path), and then expanded towards the output image (***Fig. 6***, expanding path). Each subsequent Inception module in the DeepIFC model contained two times the number of filters contained by the preceding module, up to 512 filters. In addition, there are skip connections which connect the layers of the same spatial dimension in the contracting and expanding path. Max-pooling layers are used to reduce the input dimensionality before the model bottleneck, and then upsampling layers restore the original dimensionality to generate the fluorescent image. In addition to the fluorescent image, a set of 128 features can be extracted for each cell from the model’s bottleneck layer. For UMAP visualization, features for all marker channels are concatenated to form feature files in the shape of n_cells x 128, so all markers can be visualized in the same plot. Each DeepIFC model generates a single fluorescent image for each instance in the data. Thus, a separate DeepIFC model is trained for each fluorescent channel in the input data.

**Figure 6.**
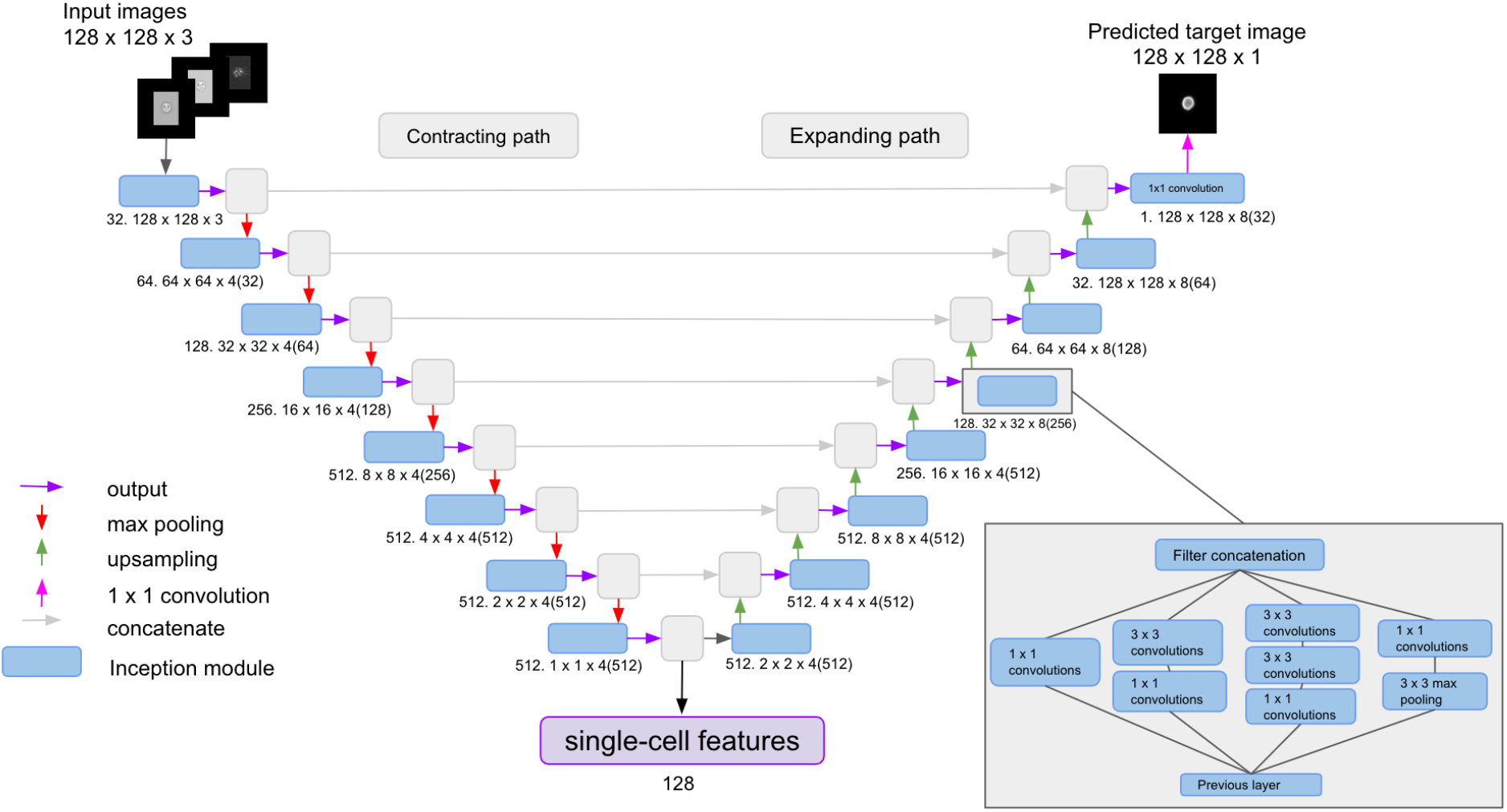
DeepIFC model architecture is based on Inception U-Net (Cahall et al. 2019). The model inputs two brightfield and one darkfield images, and generates a predicted fluorescent image. The model consists of contracting (encoding) and expanding (decoding) paths, with a bottleneck layer in the middle outputting single-cell features. Number of filters, and input dimensionality are indicated under each module. Bottom right corner: architecture of an Inception module (Szegedy et al. 2015), consisting of convolutional and max-pooling layers.

DeepIFC converts each input CIF image to Hierarchical Data Format (HDF) using the Cifconvert tool (Lippeveld *et al*. 2020), and extends images to 128×128 size by padding edges with zero values. To normalize backgrounds in fluorescent images, each image *Y* is transformed with

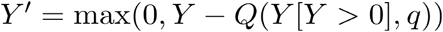

where *Q* is the quantile function and *q* was set to 0.6 in our study, to obtain images similar to those produced by IDEAS*®* software (Luminex corporation, Austin, Texas, US) (***Supplementary Fig. 1, 2***).

### Isolation and staining of the cells

Human peripheral blood mononuclear cells (PBMC) were extracted from buffy coats of blood samples from six voluntary anonymous blood donors (Finnish Red Cross Blood Service) using Ficoll-Pague™ Plus (GE Healthcare Life Sciences) density gradient centrifugation according to manufacturer’s recommendations. PBMCs were frozen to be stained later.

Thawed PBMCs, 10^6^ cells in 50 µl of staining buffer (containing 0,5 % human serum albumin - 2 mM EDTA in PBS), were stained with a panel of covalently linked fluorescent antibodies designed to identify main mononuclear cell types (PBMC panel); CD45-FITC (leukocyte marker; Biolegend), CD14-PE-Dazzle (monocyte; Biolegend), CD19-BV510 (B cell; Biolegend), and CD8-BV605 (cytotoxic T cell; Biolegend), CD3-PE (T cell; eBioscience), CD56-APC (natural killer or natural killer T cell; 1^st^ three samples eBioscience, next four samples Miltenyi Biotec) and 7-aminoactinomycin D (7-AAD, dead/damaged cell; BD Biosciences) **(*Supplementary Table 4***). Cell samples were also stained with corresponding isotype control antibodies. To reduce the background staining, cells were treated with BD Fc Block (BD Biosciences), following the manufacturer’s recommendation.

### Imaging flow cytometry of peripheral blood mononuclear cells

Seven PBMC samples of six blood donors (***Supplementary Table 5***) were imaged with a 12 channel Amnis® ImageStream®^X^ Mark II imaging flow cytometer (ISX) (Luminex) to capture images of mononuclear cells (MNC). One of two samples (NK11B) obtained from the same donor was discarded from the experiment as it contained a high amount of cells positive for the damaged/dead cell marker 7-AAD (54%) (***Supplementary Table 1***), possibly due to an error in freezing and unfreezing the sample. Images were acquired at 60× magnification with low flow rate/high sensitivity (40 mm/s, core 7 µm), pixel size 0.33×0.33 µm^2^ and depth of field 2.5 µm. During the experiment, excitation lasers 405 nm (intensity 120 mW), 488 nm (intensity 145 mW) and 642 nm (intensity 150 mW) were used. Fluorescent signals were gained using channels Ch02-Ch05, Ch08, Ch10 and Ch11. Channels Ch01 and Ch09 were used for brightfield (BF) images and channel 12 for darkfield (scattered light, SSC). Examples of images obtained from IDEAS® are shown in ***Supplementary Figure 1***.

We aimed to acquire 100,000 imaging events for each sample. Single color controls were used for compensation. Isotype controls and unlabelled cells were used to determine the auto fluorescence and non-specific signal. The integrated software INSPIRE® (EMD Millipore) was used for data collection. Both uncompensated and compensated images were created from the experiments with the IDEAS® software (version 6.2). Compensated images were created by applying a compensation matrix to uncompensated image data in IDEAS®, and used in training and evaluating the DeepIFC models. In addition to images, also numerical cell features collected during the experiment (*e*.*g*., intensity of fluorescence) were extracted from IDEAS®.

We performed a gating analysis in IDEAS® software, where positive events for each surface marker were gated based on the intensity values of fluorescent signals. Altogether two types of gating hierarchies were employed (***Supplementary Fig. 4a, 1***). In the first one, cell surface markers were gated from all events (“All”). This gating hierarchy was used to compare DeepIFC and IDEAS® results. In the second, R1 gate was set based on aspect ratio intensity and area features on channel 1 (brightfield) to exclude cell debris and cell clumps before surface marker gating (***Supplementary Figure 4a, 2***). Cell types were defined as follows in all IDEAS® gating hierarchies: leukocytes (CD45+), T cells (CD45+CD3+), Cytotoxic T cells (CD45+CD3+CD8+), Natural Killer T cells (CD45+CD3+CD56+), Natural Killer (NK) cells (CD45+CD3-CD56+), B cells (CD45+CD19+), and monocytes (CD45+CD14+). Gates were set based on isotype control samples and cell populations separated on area vs. fluorescence intensity plots (***Supplementary Fig. 6*)**.

### Cell typing

In order to classify cell types using the fluorescent images generated by DeepIFC, we implemented two cell typing strategies, permissible and strict typing (**Supplementary Table 2**). In both strategies, marker positivity depends whether the average intensity of the fluorescent image exceeds the specified threshold value. Threshold values were determined for each marker by inspecting intensity histograms and fluorescent target images, most commonly set at 0.01 (**Supplementary Fig. 7**). The threshold value for the DNA-binding 7-AAD was determined to be lower than the other six surface markers (0.007), possibly due to 7-AAD binding inside the cell, instead of the surface, as well as dead cells’ ability to bind antibodies non-specifically (Tomlinson *et al*. 2013). The threshold for CD19, CD8 and CD56 was also found to differ from the rest of the markers (0.006-0.012). To perform cell typing, each input cell was processed with all seven DeepIFC models, each predicting the fluorescent image of one marker. Permissible typing of DeepIFC model predictions was used to compare to the cell type proportions reported by IDEAS*®* to allow a cell to be assigned in both “T cell” and “dead/damaged” classes, for example. On the other hand, strict cell typing was used to report classification measures which allow each cell to be in exactly one class, which is a useful measure for excluding dead or damaged cells from other populations due to their ability to bind antibodies non-specifically, possibly distorting real cell population counts (Germann *et al*. 2013, Tomlinson *et al*. 2013). The cell type fractions obtained with IDEAS® with gating strategy 1 are shown in ***Supplementary Table 1***. Three different IDEAS*®*-based gating strategies were utilized with the data. Different gating strategies were undertaken to showcase the difference between isolating debris and dead cells from the data before cell type quantification, and leaving all of the objects in the whole population in order to mimic the raw CIF dataset. DeepIFC results were compared with gating strategy 1 (***Supplementary Fig. 4a)*** and an image-based ground truth generated from the fluorescent marker channel images.

### Finding doublet events in the data

To find leukocyte cell doublets, we extracted events that exhibited high CD45 intensity values (mean intensity >0.04; *n*=8840 events, 1.7%), as such high values can be caused by the fluorescence of two cells. We then manually removed single cell events from these candidate events. A random subset (n=622) of these cells was selected and classified into cell types by visually examining marker fluorescence intensity for each cell in the events, as it was not possible to utilize the previously implemented cell typing method (**Supplementary Table 2**) for cell doublets. Jeffrey’s confidence intervals were computed for all cell types at *α*=0.05 comparing the proportion of cells observed in doublets to the proportion in all cells.

### Training models on complete and balanced datasets

To train DeepIFC models on all (complete) IFC data obtained from the six PBMC samples stained with the MNC panel (n=527,107 cells), we assigned all images from two samples into a test dataset (200,000 cells, 38%), and divided the remaining images into training (247,107 cells, 47%) and validation (80,000 cells, 15%) datasets. Seven DeepIFC models were trained with training data, one for each individual marker channel in data.

To train models on data balanced with respect to cell types, we created a series of datasets 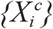 for each target cell type *c*, such that the size of each dataset 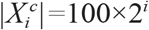, with *c* ∈ {B cell, monocyte, T cell, cytotoxic T cell, NK, NKT, dead/damaged} and *i*=2,…,6. The same two donors as in the complete dataset experiment were assigned to the test set, and other four donors to the training set. To create each dataset, 50% of cells were sampled from cells of the target cell type in the complete dataset and 50% from any other cell type. For cell types with at least 100×2^6^=6400 cells in the complete data, this resulted in a series of datasets of sizes 400, 800, 1600, 3200 and 6400 cells. For B cells and NKT cells, the largest datasets were instead 3200 and 3112 cells, respectively, due to lack of cells of the particular type in the complete data.

All DeepIFC models were trained by minimizing binary cross-entropy (BCE) of the generated fluorescent images compared to the true (target) image in a training dataset with the Adam optimizer (Kingma & Ba 2014). To train on complete data, initial learning rate 0.002 and minibatch size 20 were used. Training was stopped early after a maximum of 100 epochs or when BCE in the validation dataset did not improve for five epochs. We decided to use 8 filters in the first Inception module based on a hyperparameter search (**Supplementary Material; Supplementary Fig. 8**). To train on balanced data, initial learning rate 0.001, minibatch size 25, 500 maximum epochs, early stopping after 10 epochs, and 8 filters in the first module were used. The models were trained three times for each variation of dataset. These model iterations were tested with 1, 2, 4, 8, 16 and 32 filters, and the best balance between training time and accuracy was with 8 filters (**Supplementary Fig. 8**). The model achieving the smallest BCE in the validation dataset was retained from each experiment for further analysis. All models were trained on Nvidia Tesla V100 GPUs with 16 GB RAM (CUDA version 11.0).

### Ethical considerations

PBMCs were isolated from buffy coats that are leftover products from the standard blood donation. Voluntary blood donors have been informed about the study and they have signed the informed consent. The study has been evaluated and approved by the ethical committee of Helsinki University Hospital District HUS/1845/2019 (original statement 26.06.2019, amendments 27.11.2019 and 30.10.2020). All the samples have been treated anonymously.

## REFERENCES

Betters, D. M. (2015). Use of flow cytometry in clinical practice. Journal of the advanced practitioner in oncology, 6(5), 435.

Blasi, T., Hennig, H., Summers, H. D., Theis, F. J., Cerveira, J., Patterson, J. O., … & Rees, P. (2016). Label-free cell cycle analysis for high-throughput imaging flow cytometry. Nature communications, 7(1), 1–9.

Burel, J. G., Pomaznoy, M., Arlehamn, C. S. L., Weiskopf, D., da Silva Antunes, R., Jung, Y., … & Peters, B. (2019). Circulating T cell-monocyte complexes are markers of immune perturbations. Elife, 8, e46045.

Cahall, D. E., Rasool, G., Bouaynaya, N. C., & Fathallah-Shaykh, H. M. (2019). Inception modules enhance brain tumor segmentation. Frontiers in computational neuroscience, 13, 44.

Christiansen, E. M., Yang, S. J., Ando, D. M., Javaherian, A., Skibinski, G., Lipnick, S., … & Finkbeiner, S. (2018). In silico labeling: predicting fluorescent labels in unlabeled images. Cell, 173(3), 792–803.

Doan, M., Vorobjev, I., Rees, P., Filby, A., Wolkenhauer, O., Goldfeld, A. E., … & Hennig, H. (2018). Diagnostic potential of imaging flow cytometry. Trends in biotechnology, 36(7), 649–652.

Doan, M., Case, M., Masic, D., Hennig, H., McQuin, C., Caicedo, J., … & Irving, J. (2020). Label-free leukemia monitoring by computer vision. Cytometry Part A, 97(4), 407–414.

Doan, M., Sebastian, J. A., Caicedo, J. C., Siegert, S., Roch, A., Turner, T. R., … & Carpenter, A. E. (2020). Objective assessment of stored blood quality by deep learning. Proceedings of the National Academy of Sciences, 117(35), 21381–21390.

Eulenberg, P., Köhler, N., Blasi, T., Filby, A., Carpenter, A. E., Rees, P., … & Wolf, F. A. (2017). Reconstructing cell cycle and disease progression using deep learning. Nature communications, 8(1), 1–6.

George, T. C., Basiji, D. A., Hall, B. E., Lynch, D. H., Ortyn, W. E., Perry, D. J., … & Morrissey, P. J. (2004). Distinguishing modes of cell death using the ImageStream® multispectral imaging flow cytometer. Cytometry Part A: the journal of the International Society for Analytical Cytology, 59(2), 237–245.

Germann A, et al. (2013). Temperature fluctuations during deep temperature cryopreservation reduce PBMC recovery, viability and T-cell function. Cryobiology. 2013 Oct;67(2):193–200. doi: 10.1016/j.cryobiol.2013.06.012. Epub 2013 Jul 9. PMID: 23850825.

Golden, J. B., Groft, S. G., Squeri, M. V., Debanne, S. M., Ward, N. L., McCormick, T. S., & Cooper, K. D. (2015). Chronic psoriatic skin inflammation leads to increased monocyte adhesion and aggregation. The Journal of Immunology, 195(5), 2006–2018.

Haridas, V., Ranjbar, S., Vorobjev, I. A., Goldfeld, A. E., & Barteneva, N. S. (2017). Imaging flow cytometry analysis of intracellular pathogens. Methods, 112, 91–104.

Hennig, H., Rees, P., Blasi, T., Kamentsky, L., Hung, J., Dao, D., … & Filby, A. (2017). An open-source solution for advanced imaging flow cytometry data analysis using machine learning. Methods, 112, 201–210.

Hu, Z., Tang, A., Singh, J., Bhattacharya, S., & Butte, A. J. (2020). A robust and interpretable end-to-end deep learning model for cytometry data. Proceedings of the National Academy of Sciences, 117(35), 21373–21380.

Icha, J., Weber, M., Waters, J. C., & Norden, C. (2017). Phototoxicity in live fluorescence microscopy, and how to avoid it. BioEssays, 39(8), 1700003.

Kingma, D. P., & Ba, J. (2014). Adam: A method for stochastic optimization. arXiv preprint 1412.6980.

Kobayashi, H., Lei, C., Wu, Y., Huang, C. J., Yasumoto, A., Jona, M., … & Goda, K. (2019). Intelligent whole-blood imaging flow cytometry for simple, rapid, and cost-effective drug-susceptibility testing of leukemia. Lab on a Chip, 19(16), 2688–2698.

Larochelle, H., Erhan, D., & Bengio, Y. (2008, July). Zero-data learning of new tasks. In AAAI (Vol. 1, No. 2, p. 3).

Li, Y., Mahjoubfar, A., Chen, C. L., Niazi, K. R., Pei, L., & Jalali, B. (2019). Deep cytometry: deep learning with real-time inference in cell sorting and flow cytometry. Scientific reports, 9(1), 1–12.

Lippeveld, M., Knill, C., Ladlow, E., Fuller, A., Michaelis, L. J., Saeys, Y., … & Peralta, D. (2020). Classification of human white blood cells using machine learning for stain-free imaging flow cytometry. Cytometry Part A, 97(3), 308–319.

Luo, S., Nguyen, K. T., Nguyen, B. T., Feng, S., Shi, Y., Elsayed, A., … & Liu, A. Q. (2021). Deep learning-enabled imaging flow cytometry for high-speed Cryptosporidium and Giardia detection. Cytometry Part A, 99(11), 1123–1133.

Luo, S., Shi, Y., Chin, L. K., Hutchinson, P. E., Zhang, Y., Chierchia, G., … & Liu, A. Q. (2021). Machine-Learning-Assisted Intelligent Imaging Flow Cytometry: A Review. Advanced Intelligent Systems, 3(11), 2100073.

Matsuoka, Y., Nakatsuka, R., & Fujioka, T. (2021). Automatic discrimination of human hematopoietic tumor cell lines using a combination of imaging flow cytometry and convolutional neural network. Human Cell, 1-4.

McInnes, L., Healy, J., & Melville, J. (2018). Umap: Uniform manifold approximation and projection for dimension reduction. arXiv preprint 1802.03426.

Nassar, M., Doan, M., Filby, A., Wolkenhauer, O., Fogg, D. K., Piasecka, J., … & Hennig, H. (2019). Label-free identification of white blood cells using machine learning. Cytometry Part A, 95(8), 836–842.

Nguyen, P., Chien, S., Dai, J., Monnat Jr, R. J., Becker, P. S., & Kueh, H. Y. (2021). Unsupervised discovery of dynamic cell phenotypic states from transmitted light movies. PLoS computational biology, 17(12), e1009626.

Otesteanu, C. F., Ugrinic, M., Holzner, G., Chang, Y. T., Fassnacht, C., Guenova, E., … & Claassen, M. (2021). A weakly supervised deep learning approach for label-free imaging flow-cytometry-based blood diagnostics. Cell reports methods, 1(6), 100094.

Ounkomol, C., Seshamani, S., Maleckar, M. M., Collman, F., & Johnson, G. R. (2018). Label-free prediction of three-dimensional fluorescence images from transmitted-light microscopy. Nature methods, 15(11), 917–920.

Ronneberger, O., Fischer, P., & Brox, T. (2015, October). U-net: Convolutional networks for biomedical image segmentation. In International Conference on Medical image computing and computer-assisted intervention (pp. 234–241). Springer, Cham.

Shi, W., Jiang, F., & Zhao, D. (2017, September). Single image super-resolution with dilated convolution based multi-scale information learning inception module. In 2017 IEEE International Conference on Image Processing (ICIP) (pp. 977-981). IEEE.

Szegedy, C., Vanhoucke, V., Ioffe, S., Shlens, J., & Wojna, Z. (2016). Rethinking the inception architecture for computer vision. In Proceedings of the IEEE conference on computer vision and pattern recognition (pp. 2818–2826).

Tang, R., Zhang, Z., Chen, X., Waller, L., Zhang, A. C., Chen, J., … & Lo, Y. H. (2020). 3D side-scattering imaging flow cytometer and convolutional neural network for label-free cell analysis. APL Photonics, 5(12), 126105.

Tomlinson, M. J., Tomlinson, S., Yang, X. B., & Kirkham, J. (2013). Cell separation: Terminology and practical considerations. Journal of tissue engineering, 4, 2041731412472690.

Wang, Y., Kartasalo, K., Weitz, P., Acs, B., Valkonen, M., Larsson, C., … & Rantalainen, M. (2021). Predicting molecular phenotypes from histopathology images: A transcriptome-wide expression–morphology analysis in breast cancer. Cancer Research, 81(19), 5115–5126.

Wang, H., Rivenson, Y., Jin, Y., Wei, Z., Gao, R., Günaydın, H., … & Ozcan, A. (2019). Deep learning enables cross-modality super-resolution in fluorescence microscopy. Nature methods, 16(1), 103–110.

Wieslander, H., Gupta, A., Bergman, E., Hallström, E., & Harrison, P. J. (2021). Learning to see colours: Biologically relevant virtual staining for adipocyte cell images. PloS one, 16(10), e0258546.

