## Supplementary information for "DeepIFC: virtual fluorescent labeling of blood cells in imaging flow cytometry data with deep learning"

### SUPPLEMENTARY MATERIAL

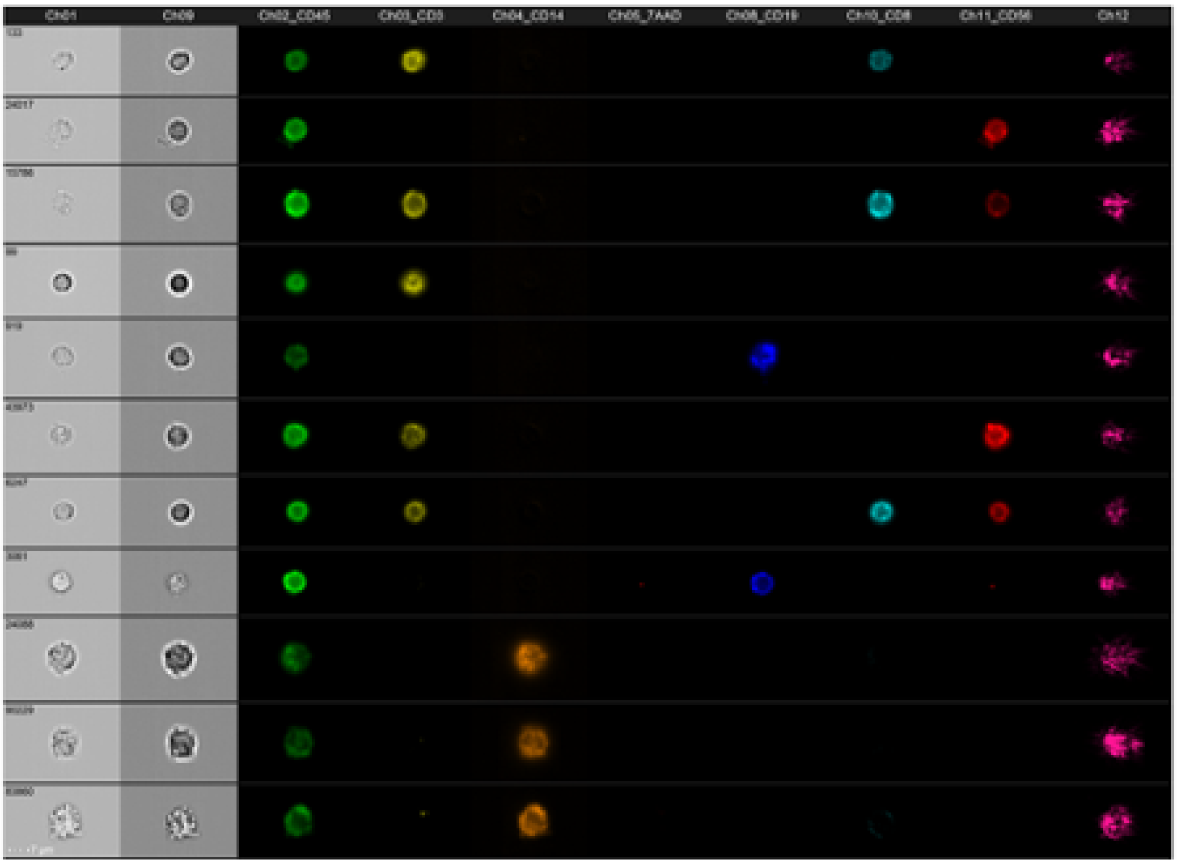

***Supplementary Figure 1***. *Example images from IDEAS® software measured in sample NK17. Each row corresponds to a single cell and each column to a channel of the imaging flow cytometer. Columns from*

*left to right: Ch01 brightfield 1, Ch09 brightfield 2, Ch02 CD45, Ch03 CD3, Ch04 CD14, CD05 7-AAD, Ch08 CD19, Ch10 CD8, Ch11 CD56, Ch12 darkfield.*

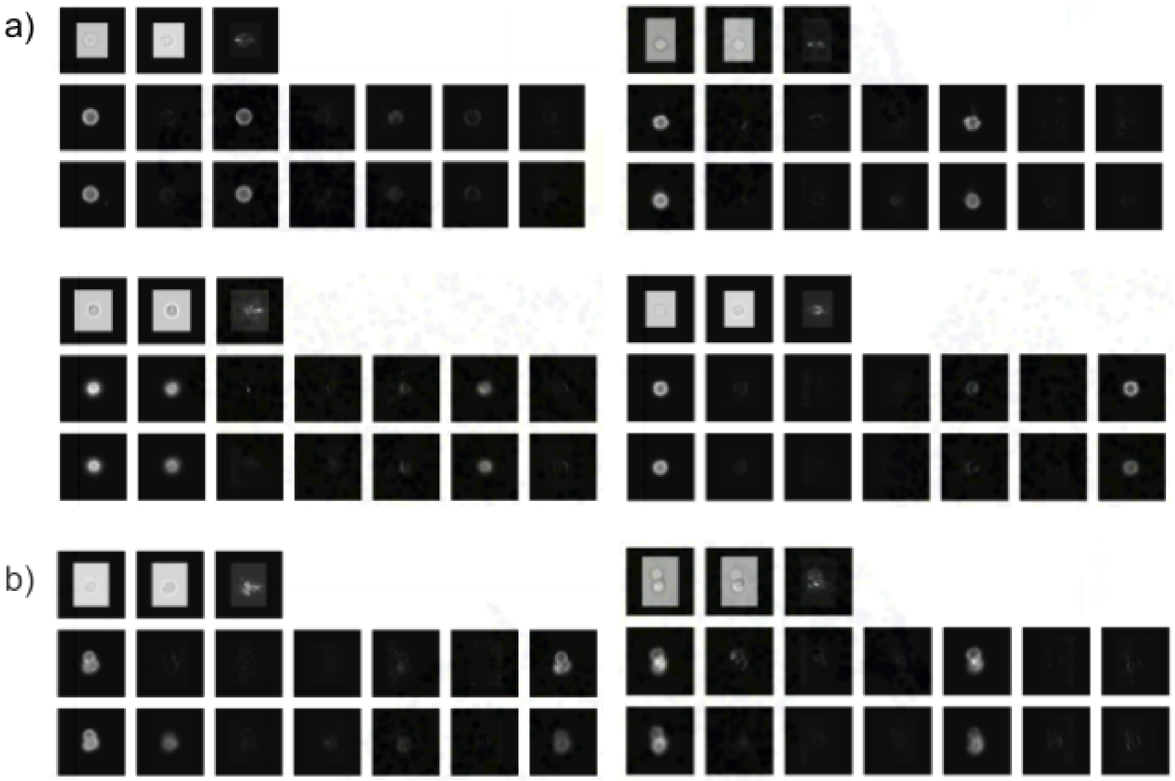


***Supplementary Figure 2.*** *Examples of cell images measured with an imaging flow cytometer and fluorescent channel images generated by the DeepIFC model. For each cell, the top, middle and bottom row correspond to brightfield and darkfield images (n=3), measured fluorescent images (n=7) and generated fluorescent images (n=7), respectively. Top row channels are, from left to right, brightfield 1, brightfield 2 and darkfield. Middle and bottom row channels are CD45, CD3, CD14, 7-AAD, CD19, CD8 and CD56. Screenshots from the DeepIFC interactive tool.* ***a)*** *Successful DeepIFC predictions: monocyte (top left), B cell (top right), cytotoxic T cell (bottom left) and NK cell (bottom right).* ***b)*** *Two examples of multiple cells in the same image: a triplet of NK cells (upper) and a triplet of B cells (lower).*

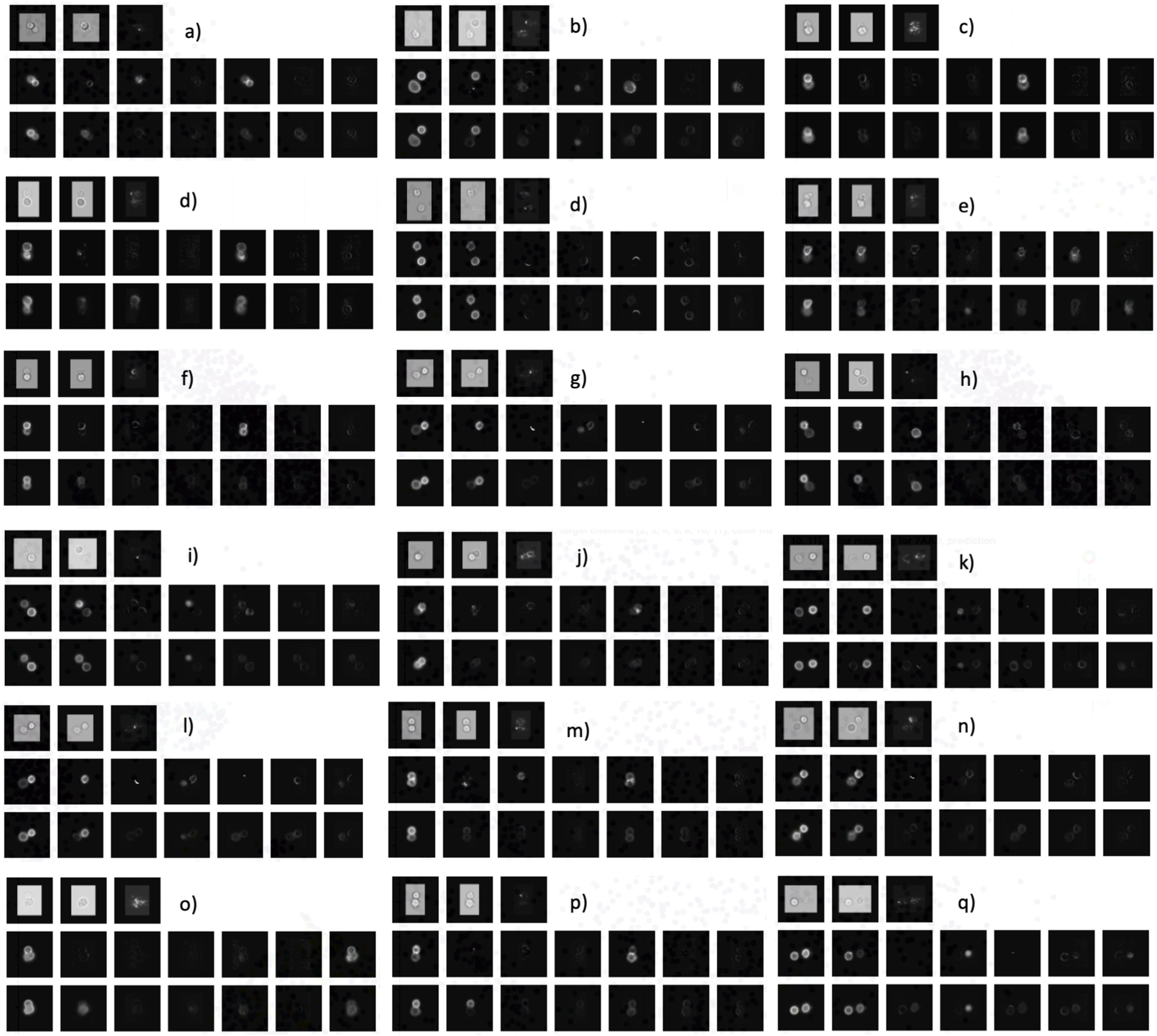


***Supplementary Figure 3.*** *Examples of doublet events (n=18). Top row channels are, from left to right, brightfield 1, brightfield 2 and darkfield. Middle and bottom row channels are CD45, CD3, CD14, 7-AAD, CD19, CD8 and CD56. The cell types for target images and DeepIFC generated predictions are as follows (as target-predicted pairs, strict typing):* ***a)*** *B, T,* ***b)*** *damaged/dead, damaged/dead,* ***c)*** *B, B,* ***d)*** *B, B,* ***e)*** *cytotoxic T, NKT,* ***f)*** *B, WBC,* ***g)*** *damaged/dead, NKT,* ***h)*** *NKT, T,* ***i)*** *damaged/dead, damaged/dead,* ***j)*** *B, WBC,* ***k)*** *damaged/dead, damaged/dead,* ***l)*** *damaged/dead, NKT,* ***m)*** *B, WBC,* ***n)*** *damaged/dead, cytotoxic T,* ***o)*** *NK, NKT,* ***p)*** *B, T,* ***q)*** *damaged/dead, damaged/dead.*

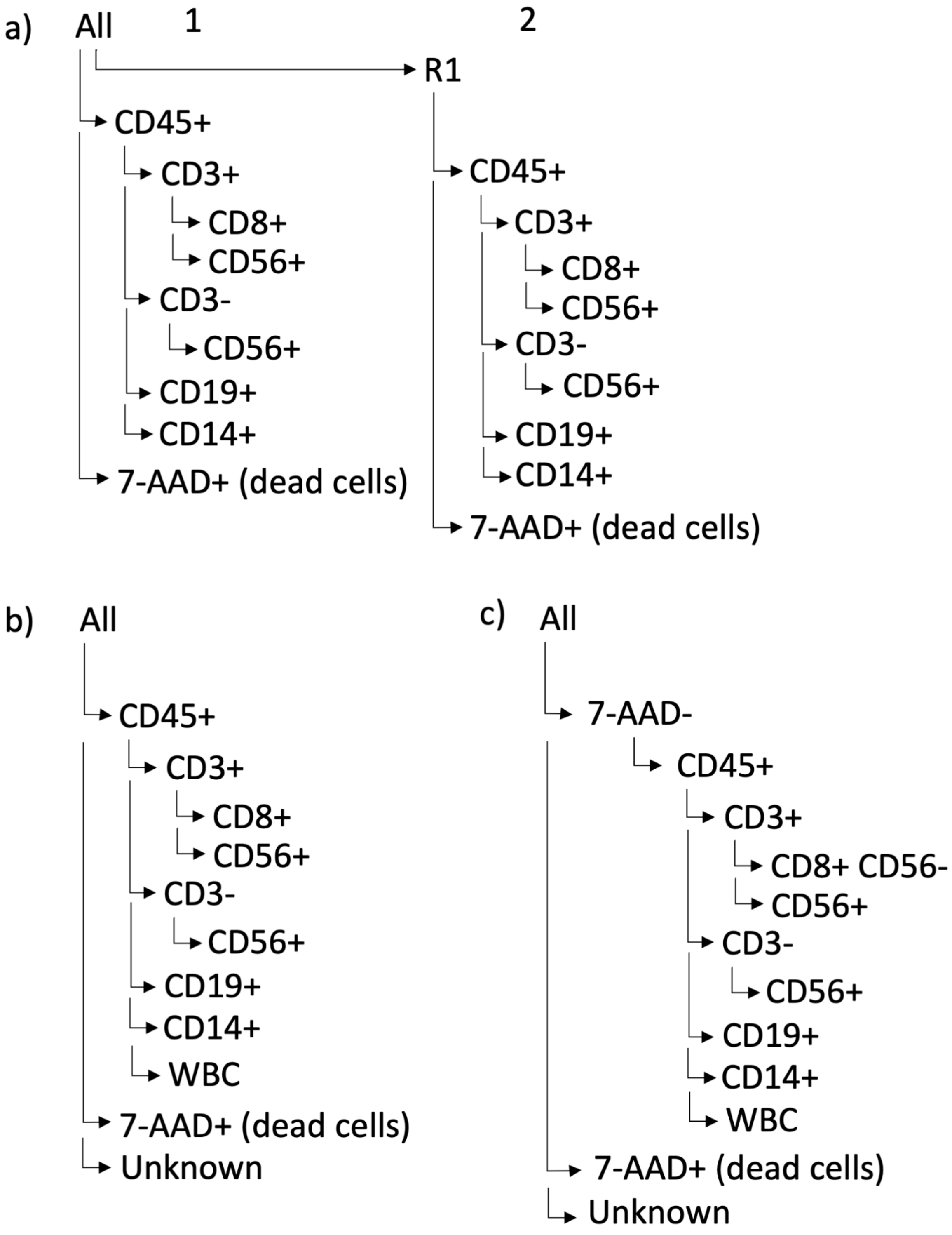


***Supplementary Figure 4.*** *Different gating strategies used in the study.* ***a)*** *IDEAS® gating hierarchy,* ***b)*** *permissible cell typing which emulates IDEAS® strategy 1,* ***c)*** *strict cell typing used to evaluate DeepIFC prediction performance.*

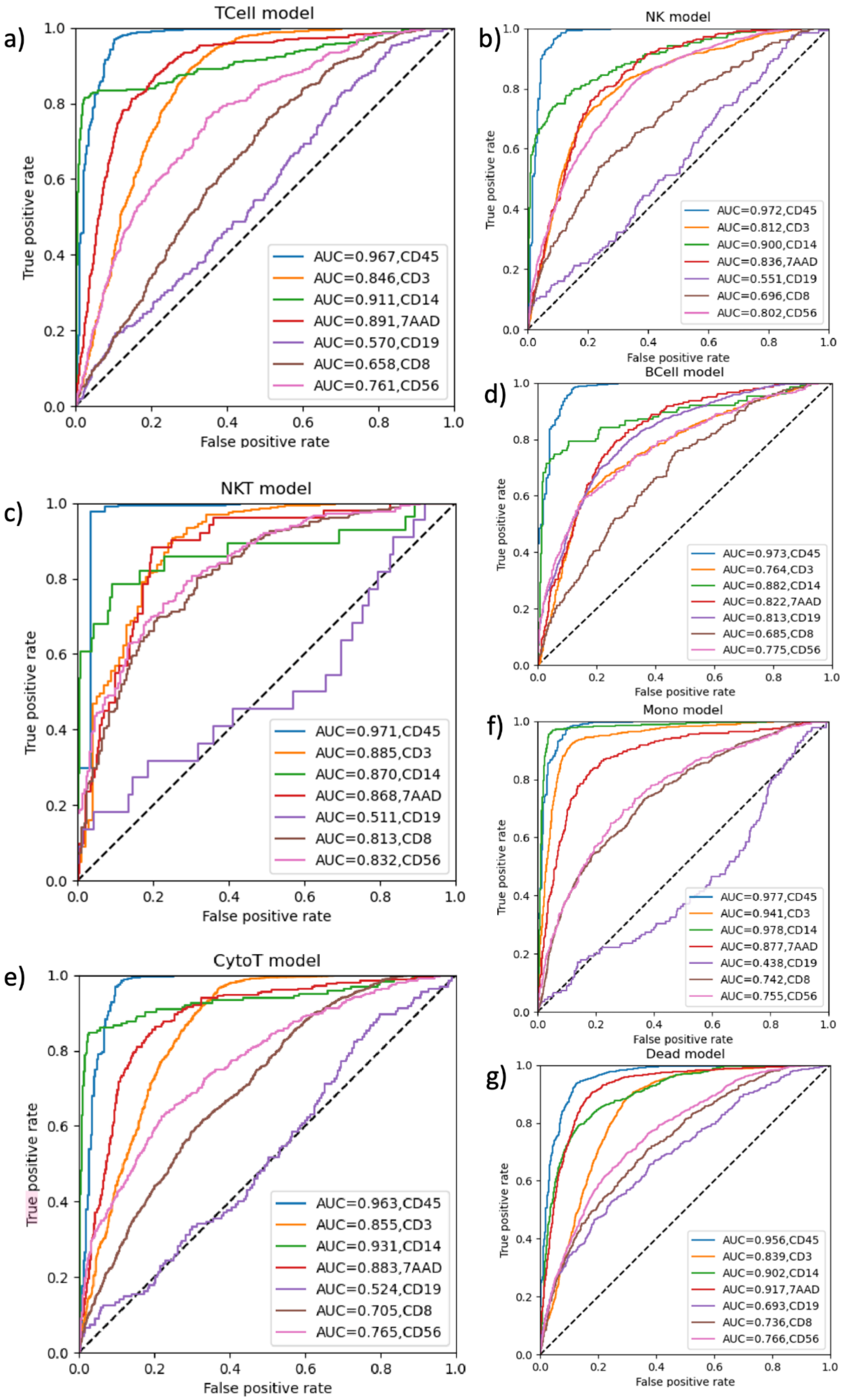
a)
TCell model
True positive rate
False positive rate
AUC=0.967,CD45
AUC=0.846,CD3
AUC=0.911,CD14
AUC=0.891,7AAD
AUC=0.570,CD19
AUC=0.658,CD8
AUC=0.761,CD56
b)
NK model
True positive rate
False positive rate
AUC=0.972,CD45
AUC=0.812,CD3
AUC=0.900,CD14
AUC=0.836,7AAD
AUC=0.551,CD19
AUC=0.696,CD8
AUC=0.802,CD56
c)
NKT model
True positive rate
False positive rate
AUC=0.971,CD45
AUC=0.885,CD3
AUC=0.870,CD14
AUC=0.868,7AAD
AUC=0.511,CD19
AUC=0.813,CD8
AUC=0.832,CD56
d)
BCell model
True positive rate
False positive rate
AUC=0.973,CD45
AUC=0.764,CD3
AUC=0.882,CD14
AUC=0.822,7AAD
AUC=0.813,CD19
AUC=0.685,CD8
AUC=0.775,CD56
e)
CytoT model
True positive rate
False positive rate
AUC=0.963,CD45
AUC=0.855,CD3
AUC=0.931,CD14
AUC=0.883,7AAD
AUC=0.524,CD19
AUC=0.705,CD8
AUC=0.765,CD56
f)
Mono model
True positive rate
False positive rate
AUC=0.977,CD45
AUC=0.941,CD3
AUC=0.978,CD14
AUC=0.877,7AAD
AUC=0.438,CD19
AUC=0.742,CD8
AUC=0.755,CD56
g)
Dead model
True positive rate
False positive rate
AUC=0.956,CD45
AUC=0.839,CD3
AUC=0.902,CD14
AUC=0.917,7AAD
AUC=0.693,CD19
AUC=0.736,CD8
AUC=0.766,CD56

***Supplementary Figure 5.*** *ROC curves for best DeepIFC models trained with balanced data. Each of the models was taught with balanced data sets containing 50% a specific cell type and 50% any other cell type. Models trained for specific cell types, from left to right, top to bottom: a) T cell, b) natural killer cell, c) natural killer T cell, d) B cell, e) cytotoxic T cell, f) monocyte and g) damaged or dead.*

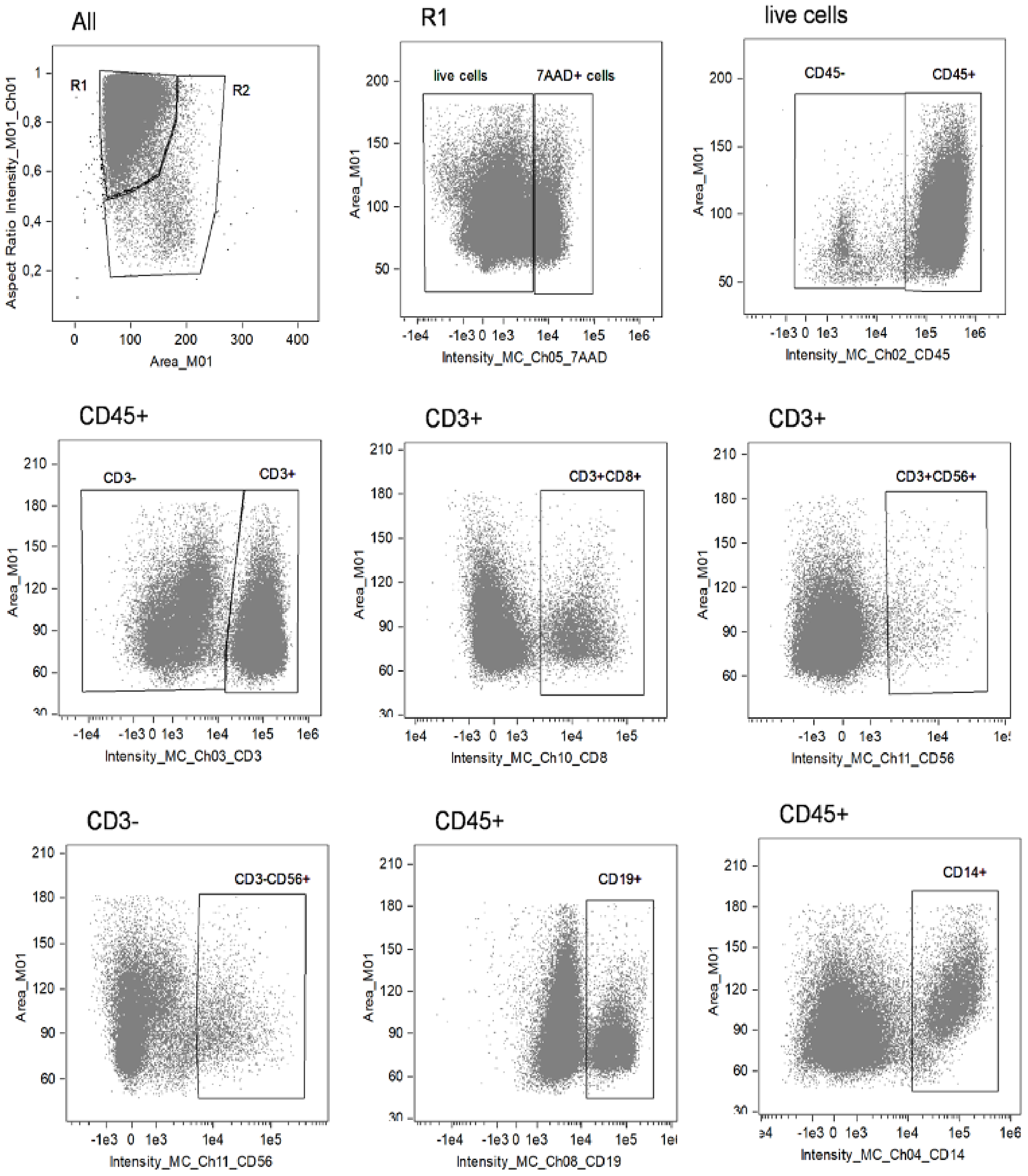


***Supplementary Figure 6.*** *Gating performed in IDEAS® to determine marker positivity and cell type counts.*

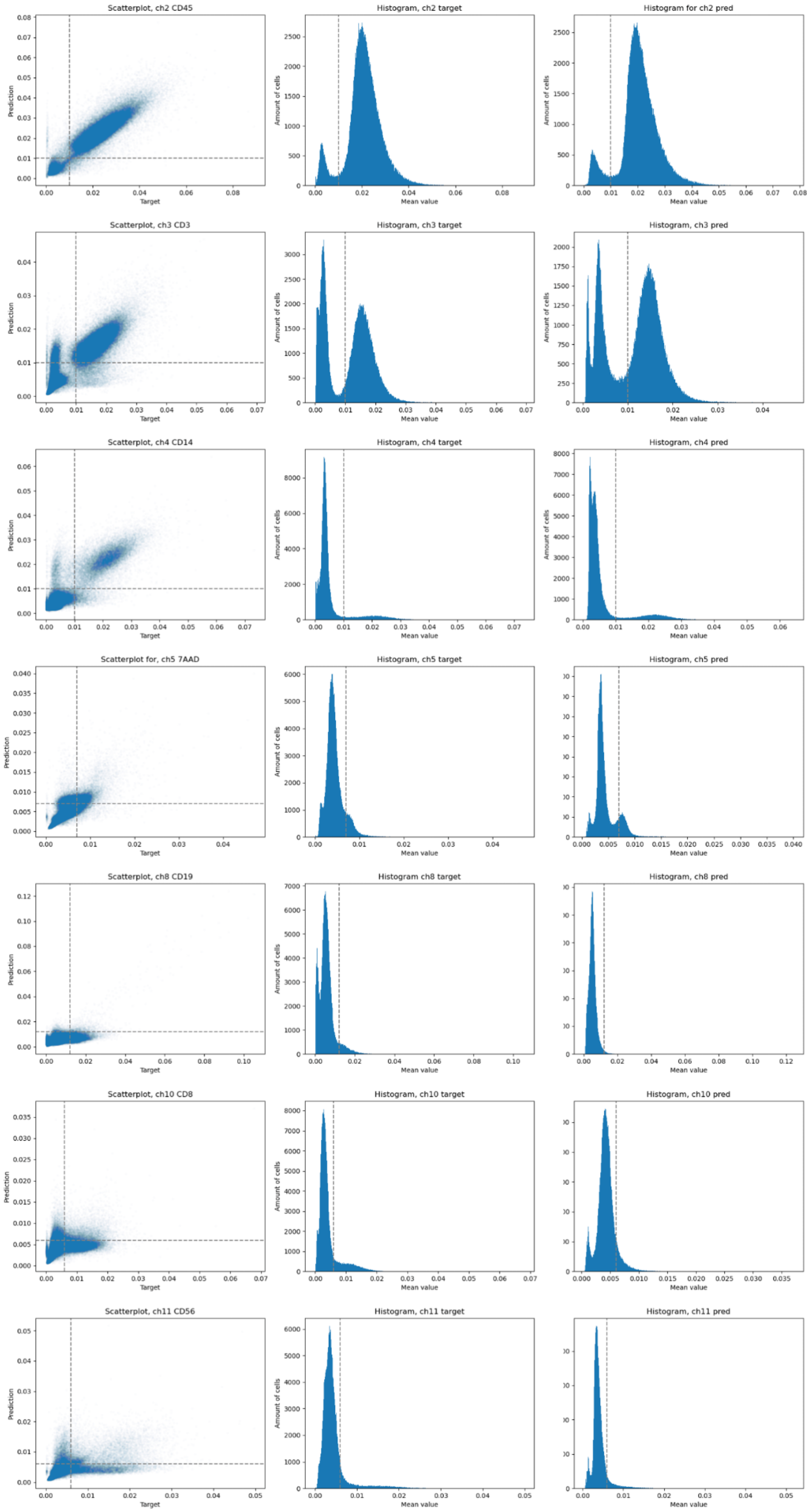
Scatterplot, ch2 CD45
Histogram, ch2 target
Histogram for ch2 pred
Scatterplot, ch3 CD3
Histogram, ch3 target
Histogram, ch3 pred
Scatterplot, ch4 CD14
Histogram, ch4 target
Histogram, ch4 pred
Scatterplot for, ch5 7AAD
Histogram, ch5 target
Histogram, ch5 pred
Scatterplot, ch8 CD19
Histogram ch8 target
Histogram, ch8 pred
Scatterplot, ch10 CD8
Histogram, ch10 target
Histogram, ch10 pred
Scatterplot, ch11 CD56
Histogram, ch11 target
Histogram, ch11 pred
Prediction
Target
Amount of cells
Mean value

***Supplementary Figure 7.*** *Distributions of observed and predicted fluorescent label intensities (n=200,000 events, test dataset, complete PBMC data). Rows: distributions of labels CD45, CD3, CD14, 7AAD, CD19, CD8 and CD56. Left column: scatterplots of observed (X-axis) vs predicted mean intensity (Y-axis). Horizontal and vertical lines denote the thresholds used to assign marker positivity. An example of histograms for the means of a 200,000 test set. Middle and right columns: distribution of predicted and observed intensities, respectively. Vertical lines denote the marker positivity threshold.*

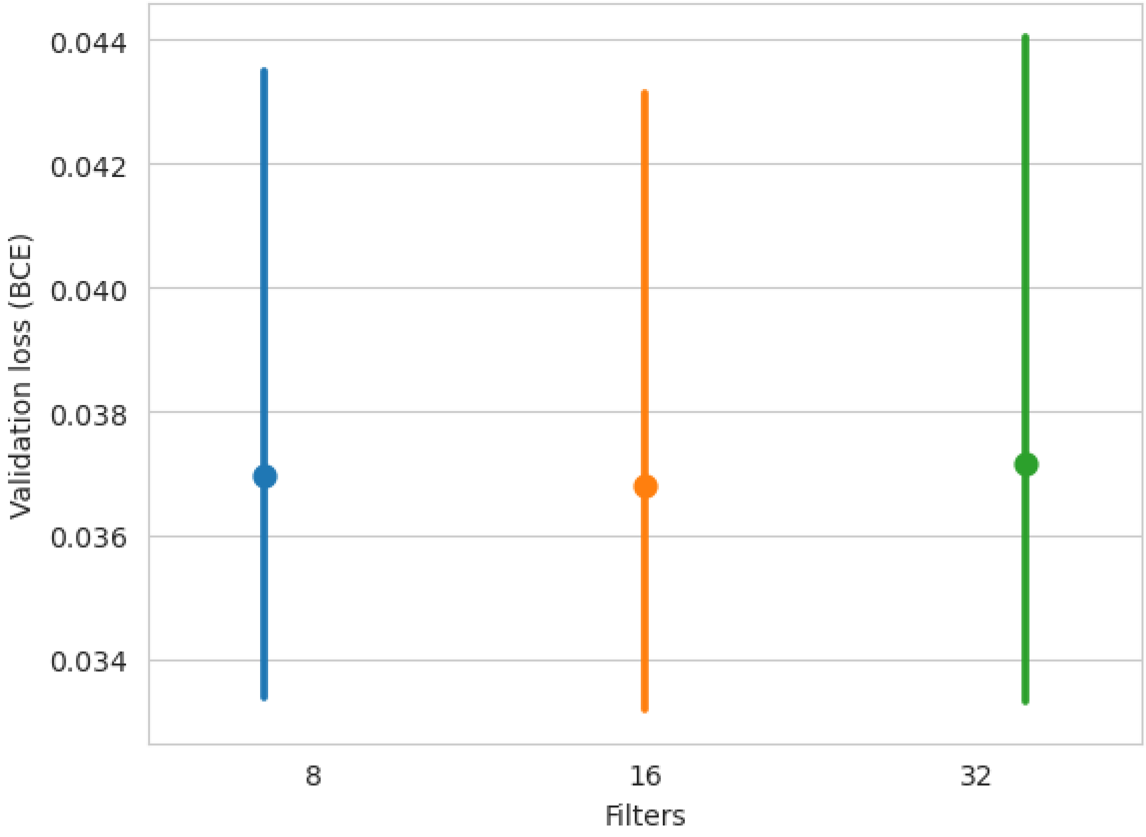


***Supplementary Figure 8.*** *Comparison between 8, 16 and 32 filters when training a DeepIFC model for marker channel 2 (CD45), showing the error bars and mean validation loss for each model. The sample used for hyperparameter testing is from donor NK11A (**Supplementary Table 5**). No difference in the best validation loss reached by the different models was observed.*

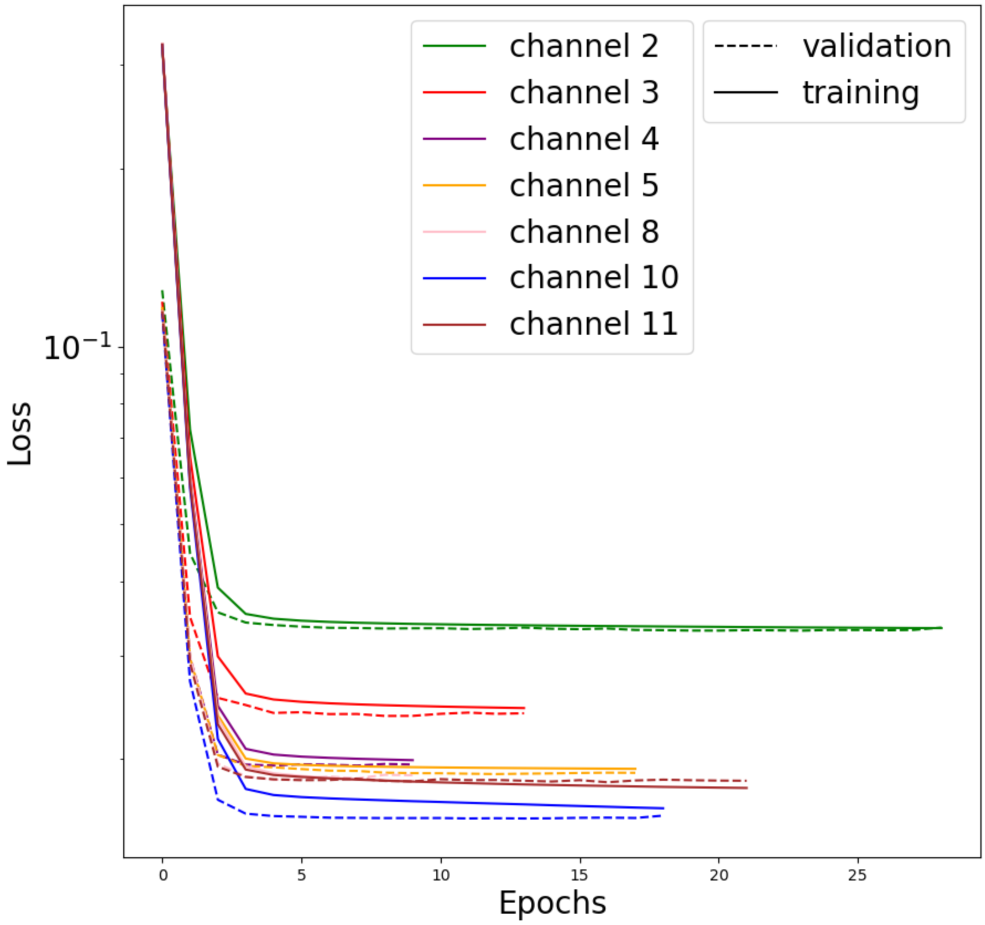


***Supplementary Figure 9.*** *The validation loss curve of each complete data model during training for each channel.*

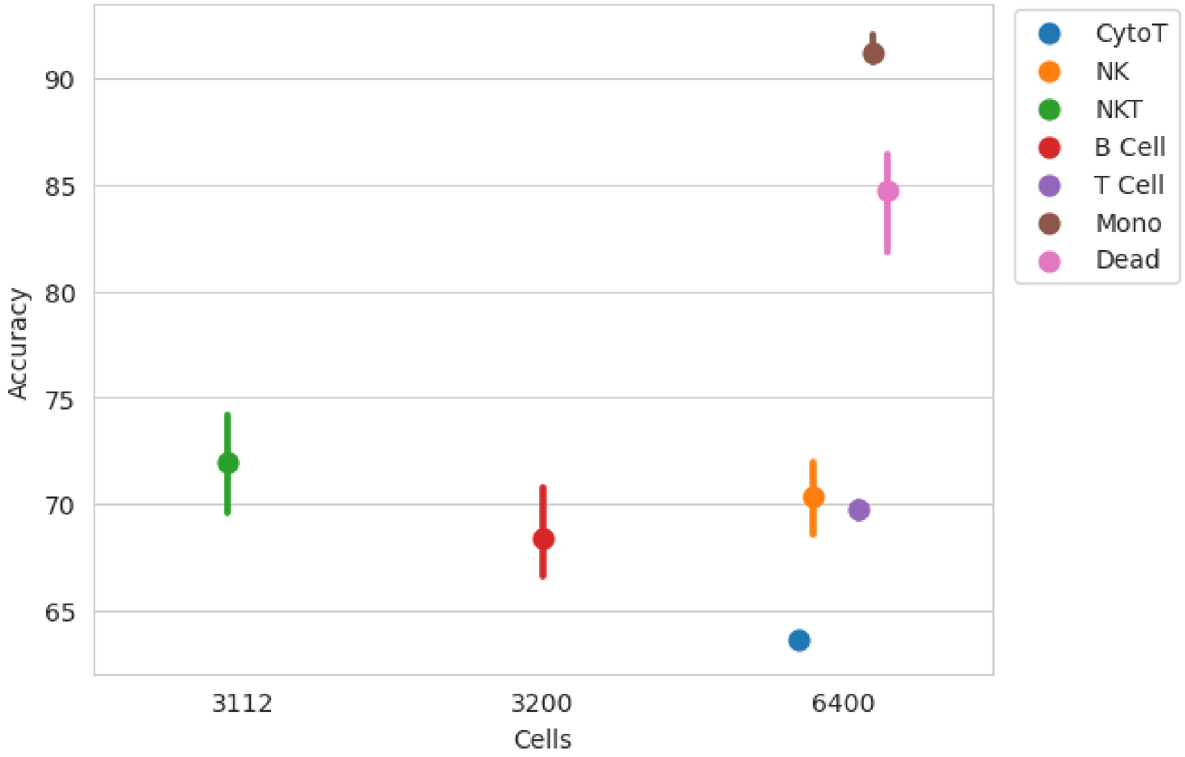


***Supplementary Figure 10.*** *Accuracies (Y-axis) of models trained on balanced data with respect to the number of cells (X-axis).*

**Supplementary Table 1**. *Cell type fractions per sample obtained from IDEAS® with gating strategy 1 (**Supplementary Fig. 4a**) including dead cells in the analysis, compared with cell counts derived from DeepIFC generated fluorescent images and observed images ("Ground truth").*

| | | Dead / Damaged, % | | | T cell, % | | | B cell, % | | | NK, % | | | NKT, % | | | Monocyte, % | | | Cytotoxic T cell, % | | |
|---|---|---|---|---|---|---|---|---|---|---|---|---|---|---|---|---|---|---|---|---|---|---|
| Sample | Cell count, all | IDEAS | DeepIFC | Ground truth | IDEAS | DeepIFC | Ground truth | IDEAS | DeepIFC | Ground truth | IDEAS | DeepIFC | Ground truth | IDEAS | DeepIFC | Ground truth | IDEAS | DeepIFC | Ground truth | IDEAS | DeepIFC | Ground truth |
| NK10 | 100000 | 16.72 | 18.25 | 12.42 | 64.21 | 65.72 | 65.73 | 13.75 | 0.41 | 7.23 | 5.13 | 4.97 | 7.73 | 7.28 | 2.84 | 3.34 | 10.59 | 8.67 | 8.43 | 7.58 | 4.57 | 5.17 |
| NK11A | 99024 | 42.01 | 33.10 | 36.40 | 63.33 | 56.97 | 60.70 | 9.59 | 0.85 | 5.05 | 7.11 | 3.73 | 4.70 | 11.85 | 1.78 | 2.01 | 11.67 | 8.21 | 9.46 | 28.61 | 5.87 | 19.0 |
| NK09 | 100000 | 17.68 | 18.35 | 16.12 | 52.57 | 51.67 | 53.44 | 5.36 | 1.04 | 3.86 | 6.04 | 15.65 | 16.56 | 10.17 | 9.47 | 9.43 | 16.75 | 18.34 | 15.35 | 14.36 | 7.17 | 10.30 |
| NK17 | 100000 | 6.55 | 8.74 | 7.18 | 57.05 | 58.47 | 57.25 | 4.61 | 0.63 | 3.31 | 7.59 | 10.50 | 14.67 | 4.04 | 4.89 | 4.87 | 12.45 | 13.08 | 9.92 | 18.20 | 6.85 | 15.62 |
| NK18 | 48060 | 18.32 | 21.38 | 12.70 | 35.69 | 33.95 | 34.83 | 3.05 | 0.32 | 1.65 | 15.14 | 13.82 | 24.0 | 7.86 | 5.06 | 5.90 | 12.58 | 11.02 | 9.49 | 8.22 | 4.21 | 4.84 |
| NK19 | 80023 | 25.26 | 24.47 | 14.28 | 55.53 | 52.83 | 52.83 | 6.20 | 0.57 | 3.64 | 8.64 | 10.68 | 13.48 | 5.51 | 3.72 | 3.09 | 8.88 | 8.76 | 6.66 | 13.49 | 5.72 | 9.09 |
| Average for all donors | **527107** | **20.0** | **20.72** | **16.52** | **64.73** | **53.27** | **54.13** | **7.09** | **0.64** | **4.12** | **8.275** | **9.9** | **13.52** | **7.785** | **4.63** | **4.77** | **12.15** | **11.35** | **9.885** | **15.08** | **5.73** | **10.67** |
| **Predicted count accuracy (DeepIFC / IDEAS)** | | 96.4 | | | 82.3 | | | 9.03 | | | 80.36 | | | 59.47 | | | 93.4 | | | 38 | | |
| **Predicted count accuracy (GT / IDEAS)** | | 82.6 | | | 83.62 | | | 58.11 | | | 36.616 | | | 61.27 | | | 81.358 | | | 70.755 | | |
| **Predicted count accuracy (DeepIFC / GT)** | | 74.576 | | | 98.4112 | | | 15.53 | | | 73.224 | | | 97.06 | | | 85.17956 | | | 53.70 | | |

**Supplementary Table 2.** *Cell typing strategies employed in DeepIFC. Plus (+) and minus (-) denote marker positivity and negativity, respectively, with +/- indicating both options are allowed for the cell type. If a cell does not match any type, it is assigned type "Unknown". **a**) In permissible cell typing, a cell be assigned multiple types. Permissible typing follows the gating strategy performed in IDEAS® with the addition of cell types white blood cell (WBC), which is only positive for the CD45 marker and negative for all others, as well as Unknown, which has an unknown combination of markers or is negative for all markers, for example a red blood cell (RBC). **b**) In strict cell typing, a cell can only be assigned a single type.*

*a) Permissible cell typing*

| PERMISSIBLE | CD45 | CD3 | CD14 | 7-AAD | CD19 | CD8 | CD56 |
|---|---|---|---|---|---|---|---|
| **White blood cell** | + | +/- | +/- | +/- | +/- | +/- | +/- |
| **T cell** | + | + | - | +/- | - | +/- | +/- |
| **Monocyte** | + | - | + | +/- | - | - | - |
| **Dead or damaged cell** | +/- | +/- | +/- | + | +/- | +/- | +/- |
| **B cell** | + | - | - | +/- | + | - | - |
| **Cytotoxic T cell** | + | + | - | +/- | - | + | - |
| **NK cell** | + | - | - | +/- | - | - | + |
| **NKT cell** | + | + | - | +/- | - | +/- | + |

*b) Strict cell typing*

| STRICT | CD45 | CD3 | CD14 | 7-AAD | CD19 | CD8 | CD56 |
|---|---|---|---|---|---|---|---|
| White blood cell | + | - | - | - | - | - | - |
| T cell | + | + | - | - | - | - | - |
| Monocyte | + | - | + | - | - | - | - |
| Dead or damaged cell | - | - | - | + | - | - | - |
| B cell | + | - | - | - | + | - | - |
| Cytotoxic T cell | + | + | - | - | - | + | - |
| NK cell | + | - | - | - | - | - | + |
| NKT cell | + | + | - | - | - | + | + |

***Supplementary Table 3.*** *The improvement of prediction and recall in balanced data over complete data. The table reports the precision and recall of the best models trained on complete data and balanced data, and the difference of the two (in percentage points, p.p.), as well as the performance of the complete model without including Unknown, Dead / Damaged and WBC cell types. The values are calculated with the strict cell typing strategy (***Supplementary Table 2***). Full confusion matrices are shown in Supplementary Table 6.*

| CELL TYPE PREDICTION (precision %; recall %) | B cell | Natural killer cell (NK) | Natural killer T cell (NKT) | Cytotoxic T cell | T cell | Monocyte | Damaged / Dead |
|---|---|---|---|---|---|---|---|
| **Complete data** | 51.7%<br>4.6% | 52.9%<br>24.8 % | 21.1%<br>23.7 % | 25.7%<br>12.8% | 73.2%<br>80.3% | 78.0%<br>90.3% | 53.5%<br>73.4% |
| **Complete data (w/o Dead/Unk/WBC)** | 67.2%<br>12.6% | 67.4%<br>44.7% | 25.9%<br>26.8% | 27.4%<br>14.1% | 77.8%<br>90.1% | 91.6%<br>98.2% | *NA* |
| **Balanced data** | 74.0%<br>54.0% | 70.8%<br>67.1 % | 62.9%<br>82.6% | 60.2%<br>67.5 % | 71.2%<br>78.5% | 90.6%<br>92.1% | 85.7%<br>87.6% |
| **Difference (with all cell types)** | **Precision**<br>+22.3 p.p.<br>**Recall**<br>+49.5 p.p. | **Precision**<br>+17.9 p.p.<br>**Recall**<br>+42.3 p.p. | **Precision**<br>+41.8 p.p.<br>**Recall**<br>+58.9 p.p. | **Precision**<br>+34.5 p.p.<br>**Recall**<br>+54.7 p.p. | **Precision**<br>-2.0 p.p.<br>**Recall**<br>-1.8 p.p. | **Precision**<br>+12.6 p.p.<br>**Recall**<br>+1.8 p.p. | **Precision**<br>+32.2 p.p.<br>**Recall**<br>+14.2 p.p. |

***Supplementary Table 4.*** *PBMC staining panel for the Amnis® ImageStream®X Mark II imaging flow cytometer used in the study. Channels 1, 9 and 12 were used as input data for the model, and channels 2, 3, 4, 5, 8, 10 and 11 as target data.*

| **Laser** | **Brightfield illumination** | **BLUE 488** | **BLUE 488** | **BLUE 488** | **BLUE 488** |
|---|---|---|---|---|---|
| **Channel** | 1 | 2 | 3 | 4 | 5 |
| **Filter** | 457/45<br>435-480 | 528/65<br>480-560 | 577/35<br>560-595 | 610/30<br>595-642 | 702/85<br>642-745 |
| **Label/BF/SSC** | Brightfield 1 | CD45-FITC | CD3-PE | CD14-PE/Dazzle | 7-AAD |

| **Laser** | **VIOLET 405** | **Brightfield illumination** | **VIOLET 405** | **RED 642** | **Darkfield 785** |
|---|---|---|---|---|---|
| **Channel** | 8 | 9 | 10 | 11 | 12 |
| **Filter** | 537/65<br>505-570 | 582/25<br>570-595 | 610-30<br>595-642 | 702/65<br>642-745 | 762/35<br>745-780 |
| **Label/BF/SSC** | CD19-BV510 | Brightfield 2 | CD8-BV605 | CD56/APC | SSC, scattered |

***Supplementary Table 5.*** *Sample donor age and sex, as well as cell counts for each cell type per sample, generated in the IDEAS® analysis.*

| **Sample** | **Sex** | **Age** | **Count, All** | **Live cells, count from All** | **Dead cells, count from All** | **Leukocytes, count from All** | **T cells, count from All** | **Cytotoxic T cells, count from All** | **NKT cells, count from All** | **NK cells, count from All** | **B cells, count from All** | **Monocytes, count from All** |
|---|---|---|---|---|---|---|---|---|---|---|---|---|
| **NK10** | F | 34 | 100000 | 79694 | 16723 | 94030 | 64207 | 7581 | 7275 | 5134 | 13751 | 10591 |
| **NK11A** | F | 36 | 99024 | 54054 | 41604 | 92039 | 62711 | 28332 | 11738 | 7040 | 9494 | 11557 |
| ***NK11B*** | *F* | *36* | *92975* | *39442* | *48767* | *83644* | *63768* | *27185* | *13339* | *5463* | *7573* | *3825* |
| **NK09** | M | 57 | 100000 | 68745 | 17679 | 83632 | 52566 | 14361 | 10168 | 6039 | 5358 | 16748 |
| **NK17** | F | 23 | 100000 | 80701 | 6547 | 84167 | 57048 | 18199 | 4039 | 7589 | 4608 | 12449 |
| **NK18** | F | 28 | 48060 | 29886 | 8805 | 34648 | 17153 | 3949 | 3778 | 7276 | 1466 | 6045 |
| **NK19** | F | 29 | 80023 | 52004 | 20213 | 66105 | 44438 | 10797 | 4410 | 6913 | 4960 | 7110 |

***Supplementary Table 6.*** *Confusion matrices for DeepIFC models trained on complete and balanced datasets. (See a separate file)*

### Supplementary Methods

#### Inception U-Net

A crucial hyperparameter choice in Inception U-Net architecture is the number of filters used in the Inception modules. We searched for an optimal choice for the number of filters by training and evaluating models with the first Inception module containing $2^k$ filters, where $k=1,...,6$. There was no considerable change in the lowest validation loss achieved as the filter size increased, so the decision was made to favor a lower amount of filters to save time in model training. The marginally best performance was obtained with 16 and $2^5=32$ filters, whereas a model with 64 filters was too large to fit in the GPU memory and could not be used (***Supplementary Fig. 8)***. Also shown is the descent of validation loss for the complete data model, for all marker channels (***Supplementary Fig. 9***), as well as the accuracy of balanced models for different sized datasets (***Supplementary Fig. 10***).
